# The impact of human activities on Australian wildlife

**DOI:** 10.1101/452409

**Authors:** Alyce Taylor-Brown, Rosie Booth, Amber Gillett, Erica Mealy, Steven Ogbourne, Adam Polkinghorne, Gabriel C. Conroy

## Abstract

Increasing human population size and the concomitant expansion of urbanisation significantly impact natural ecosystems and native fauna globally. Successful conservation management relies on precise information on the factors associated with wildlife population decline, which are challenging to acquire from natural populations. Wildlife Rehabilitation Centres (WRC) provide a rich source of this information. However, few researchers have conducted large-scale longitudinal studies, with most focussing on narrow taxonomic ranges, suggesting that WRC-associated data remains an underutilised resource, and may provide a fuller understanding of the anthropogenic threats facing native fauna.

We analysed admissions and outcomes data from a WRC in Queensland, Australia Zoo Wildlife Hospital, to determine the major factors driving admissions and morbidity of native animals in a region experiencing rapid and prolonged urban expansion.

We studied 31,626 admissions of 83 different species of native birds, reptiles, amphibians, marsupials and eutherian mammals from 2006 to 2017. While marsupial admissions were highest (41.3%), admissions increased over time for all species and exhibited seasonal variation (highest in Spring to Summer), consistent with known breeding seasons.

Causes for admission typically associated with human influenced activities were dominant and exhibited the highest mortality rates. Car strikes were the most common reason for admission (34.7%), with dog attacks (9.2%), entanglements (7.2%), and cat attacks (5.3% also high. Admissions of orphaned young and overt signs of disease were significant at 24.6% and 9.7%, respectively. Mortality rates were highest following dog attacks (72.7%) and car strikes (69.1%) and lowest in orphaned animals (22.1%).

Our results show that WRC databases offer rich opportunities for wildlife monitoring and provide quantification of the negative impacts of human activities on ecosystem stability and wildlife health. The imminent need for urgent, proactive conservation management to ameliorate the negative impacts of human activities on wildlife is clearly evident from our results.

## Introduction

There is substantive evidence to suggest that anthropogenic factors are having devastating consequences on native fauna, both in Australia [1-9] and internationally [10-26]. The stability of entire ecosystems is consistently compromised through the process of urban expansion and global population growth, which continue to increase at unprecedented rates [27]. The sustained acceleration in human population growth and resulting expansion in anthropogenic activities appear to be the primary causes of an accelerated increase in extinction rates globally [28-32].

Global population growth contributes to the destruction, modification and fragmentation of wildlife habitat, reduced genetic diversity, threats from pathogens, the spread of exotic and invasive species, air, noise and light pollution, alteration in natural hydrologic and fire regimes, and a rapidly changing climate [33-37]. The consequences of these environmental changes for most species include a reduced ability to forage, reduced prey or food availability, altered immune function, and diminished breeding success [38-45]. Changes to any of these life traits can compromise the persistence of native fauna populations in the wild.

Conception and implementation of effective conservation management strategies should be guided by a thorough understanding of the underlying causes of wildlife population decline [18, 19, 46-48]. Evaluation of longitudinal data from wildlife rehabilitation centres (WRC), including causes of admission and resultant outcomes, can be used to conduct general wildlife monitoring and investigate threats to local species [6-8, 13, 18, 19, 23, 24, 26, 47, 49-53], and may provide information about ecosystem health and stability [53, 54], quantify and delineate natural and anthropogenic elements that present potential hazards to wildlife survival.

Previous research using WRC admissions data has generally concentrated on either a single species or narrow taxonomic clusters [6-8, 15, 17, 20, 23, 50, 55, 56], with understandable foci on threatened taxa. Others have focused on particular threats, such as cat attacks, land clearing and emerging diseases [16, 21, 22, 25, 57], which have increased as human activities have encroached on wildlife habitat [26, 58].

This study takes a broader perspective, by examining a wide suite of species in South-East Queensland (QLD), Australia, including representatives from a variety of taxonomic, life history and trophic groups. The overall objective of this research was to investigate the major causes and patterns of WRC admissions and outcomes, with a sub-aim of identifying opportunities to provide targeted management solutions. The results of this longitudinal retrospective study have wide ramifications, particularly where impacts from anthropogenic processes are implicated.

## Methods

### Study site

We collated hospital records from the Australia Zoo Wildlife Hospital (AZWH) in Beerwah, Queensland. AZWH has collected data from all wildlife admissions since its opening in 2004. AZWH is located 80 km north of Brisbane, on the Sunshine Coast, which is a rapidly growing residential and tourist area, with mixed land use comprising a combination of rural, urban, peri-urban, bushland and coastal zones. The majority of AZWH admissions come from an area spanning approximately 200 km north (to Maryborough, with occasional admissions from as far north as Proserpine), 150 km west (e.g. Gatton and Kingaroy) and up to 300 km south (Lismore, New South Wales; mostly Koala admissions) (Figure 1), although admissions from central western QLD and the Northern Territory also sporadically occur.

**Figure 1:**
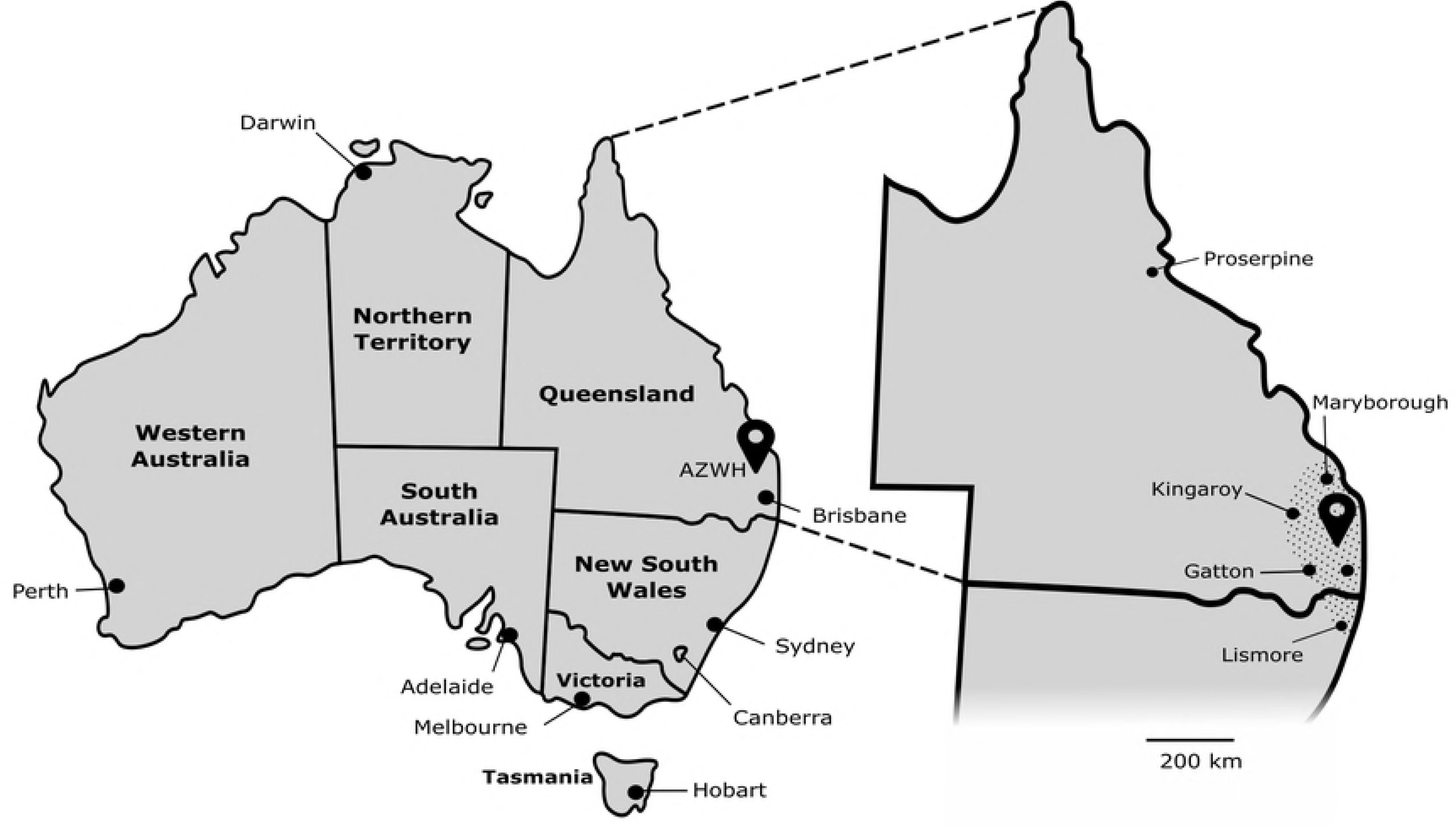
Location of Australia Zoo Wildlife Hospital (AZWH). Map of Australian states and territories, showing the location of AZWH, with a zoomed-in image of Queensland demonstrating the common admissions area of AZWH (hashed area). Scale bar is representative for the zoomed in image.

AZWH was established as a wildlife treatment facility (previously The Australian Wildlife Hospital) in March 2004. Due to a rapidly growing wildlife admission rate, a new purpose-built facility was constructed in November 2008 and is one of the largest WRCs in the world. The AZWH facilities include multiple state of the art triage assessment areas, intensive care and rehabilitation wards customised for birds, reptiles, mammals and orphaned young; radiology, laboratory, surgery and pathology facilities; and multiple large outdoor rehabilitation enclosures. It operates 24 hours a day with a team of wildlife veterinarians, vet nurses and volunteers attending to the needs of up to 8,000 wildlife admissions annually.

### Data collection

Data for 74,230 admissions between 1^st^ January 2006 and 31^st^ December 2017 were obtained from AZWH. Of these, 42,604 admission records were excluded as follows: a) data for which there were unknown, multiple, or ambiguous cause for admission (CFA) were removed from the analysis; b) admissions of animals that were dead on arrival (DOA); c) species for which there were less than 100 admissions over the time period, unless they could be suitably pooled and were a taxonomic group of interest e.g. Amphibians (see below); d) admissions of marine animals, which occupy a specific niche that we believe warrants its own detailed investigation in future studies (with the exception of the Australian pelican which had significant admission numbers from predominantly freshwater sources); e) admissions for which the outcome was not reported.

Where data on a single species was insufficient (i.e. <100 admissions) for meaningful analysis of admission and outcome trends following the exclusion criteria above, but the species was part of a larger taxonomic or ecological group of interest, we pooled these species to create a ‘multi-species group’ (Supporting File 1). Species were grouped based on either taxonomy (e.g. ‘small macropods’ are small-bodied species within the Macropodidae family, compared to Eastern grey kangaroos for example, which are larger macropods) or behaviour (e.g. raptors are a group of birds of prey that include representatives from several families) (Supporting File 1). For simplicity, taxa are referred to by their higher taxonomic groupings, termed ‘animal groups’ throughout the manuscript (i.e. avians, reptiles, amphibians, marsupial mammals and eutherian mammals; Supporting File 1).

The final dataset of 31,626 individual admissions included terrestrial and freshwater wildlife species of differing age classes, taxonomic classes and trophic groups. These data were analysed for admission and outcome trends. Where trends were assessed per season, seasons are referred to as; Summer: December, January, February; Autumn: March, April, May; Winter: June, July, August; Spring: September, October, November. CFA were listed as per their categories in the admission/accession sheets, with some CFA pooled (e.g. Bush fire and fire; Supporting file 2). Animal outcomes following admission were also grouped into either ‘positive outcome’ (release into wild or into care) or ‘mortality’ (natural death and euthanasia on welfare grounds).

Throughout the period of interest for this study, various alterations were made to the data collection methods at AZWH in response to the expansion of overall admissions and improvements in data capture methodology. Changes included; 1) intermittent updating (addition or deletion of some CFA categories) of animal admission/accession sheets; 2) restructuring of animal admission/accession sheets and redevelopment of the internal database (largely in mid-2013). Subsequently, some CFA categories were subject to change throughout the study period and may not have been clearly represented in data prior to mid-2013. To assess whether these changes might significantly alter the main findings, we performed a small subset analysis on data from 2014-2017 to evaluate any shifts in the main CFA after the changes.

### Data analysis

The aggregate data used for this study was sourced and processed through MySQL using 117 lines of SQL queries layered upon a set of 3 (112 lines total) SQL/PSM functions (Structured Query Language/ Persistent Stored Modules) [59]. Designed to maximise consistency of data, and also to allow the pooling of outcomes and species, the functions were used to filter and aggregate the raw data and to generate comma separated (csv) files. One exception was the per month/year data that was further processed using simple Java command-line application of 230 lines of code to collate the up to 3,700 data values for each of eight sheets.

The csv files were imported into Microsoft Excel and manipulated into tables of total admitted animals, causes for admission and outcomes, and grouped according to higher classification. Microsoft Excel was also used to calculate the summary statistics (totals, means and proportions), and to generate graphical outputs.

### Statistical analysis

Data were imported into IBM SPSS Statistics v24.0 [60] and reformatted where necessary. Data were assessed for distribution prior to parametric or non-parametric inferential analyses. For data with normal distribution, we performed one-way ANOVA with a Tukey Post-hoc test, and for data with non-normal distribution, we performed Kruskal-Wallis ANOVA. We used a statistical significance level of 0.05, and also performed odds-ratio analysis using the risk estimate statistic in the cross-tabs option, with a 95% confidence interval.

## Results

We studied 31,626 native animal admissions to the AZWH, a large WRC in Beerwah, Queensland, Australia, and the outcomes of those admissions, from January 2006 to December 2017. A summary of admissions over this time period is found in Table 1. A total of 83 species were included in this study, which were grouped by taxonomy, ecological niche or behavioural traits to assist analysis (Supporting File 1).

**Table 1:** Summary of admissions to AZWH from 2006 to 2017.

| Animal group | Species | Number of admissions | % of total admissions | % of animal group | % Positive outcome<sup>1</sup> | % Mortality<sup>2</sup> | Leading CFA |
| --- | --- | --- | --- | --- | --- | --- | --- |
| AVIANS | Australian Brush-turkey | 431 | 1.4% | 3.9% | 31.1% | 68.9% | HBC |
|  | Australian Magpie | 1263 | 4.0% | 11.3% | 37.1% | 62.9% | HBC |
|  | Australian Pelican | 245 | 0.8% | 2.2% | 83.3% | 16.7% | Entanglements |
|  | Fig Bird | 762 | 2.4% | 6.8% | 51.8% | 48.2% | Orphaned |
|  | Laughing Kookaburra | 1741 | 5.5% | 15.6% | 37.3% | 62.7% | HBC |
|  | Lorikeets | 2625 | 8.3% | 23.6% | 24.8% | 75.2% | HBC |
|  | Native Ducks | 948 | 3.0% | 8.5% | 67.3% | 32.7% | Orphaned |
|  | Noisy Minor | 842 | 2.7% | 7.6% | 39.1% | 60.9% | Entanglements |
|  | Raptors | 351 | 1.1% | 3.2% | 44.4% | 55.6% | HBC |
|  | Tawny Frog Mouth | 1920 | 6.1% | 17.3% | 27.1% | 72.9% | HBC |
|  |  | **11128** | **35.2%** |  | **44.3%** | **55.7%** | **HBC** |
| REPTILES | Bearded Dragon | 404 | 1.3% | 8.8% | 31.4% | 68.6% | HBC |
|  | Blue-tongued Skink | 930 | 2.9% | 20.4% | 27.6% | 72.4% | Dog attack |
|  | Carpet Python | 888 | 2.8% | 19.4% | 51.0% | 49.0% | HBC |
|  | Eastern Water Dragon | 856 | 2.7% | 18.7% | 39.3% | 60.7% | HBC |
|  | Freshwater Turtle | 646 | 2.0% | 14.1% | 52.9% | 47.1% | HBC |
|  | Green Tree Snake | 243 | 0.8% | 5.3% | 42.4% | 57.6% | Cat attack |
|  | Lace Monitor | 306 | 1.0% | 6.7% | 43.8% | 56.2% | HBC |
|  | Venomous Snakes | 295 | 0.9% | 6.5% | 52.5% | 47.5% | Cat attack |
|  |  | **4568** | **14.4%** |  | **42.6%** | **57.4%** | **HBC** |
| AMPHIBIANS | Tree Frogs | 106 | 0.3% | 100.0% | 32.1% | 67.9% | HBC |
|  |  | **106** | **0.3%** |  | **32.1%** | **67.9%** | **HBC** |
| MARSUPIALS | Bandicoots | 367 | 1.2% | 2.8% | 42.2% | 57.8% | Orphaned |
|  | Eastern Grey Kangaroo | 1165 | 3.7% | 8.9% | 30.6% | 69.4% | HBC |
|  | Feathertail Glider | 230 | 0.7% | 1.8% | 63.0% | 37.0% | Cat attack |
|  | Koala | 3590 | 11.4% | 27.5% | 44.8% | 55.2% | HBC |
|  | Large Gliders | 754 | 2.4% | 5.8% | 55.2% | 44.8% | Orphaned |
|  | Marsupial Dasyurid | 190 | 0.6% | 1.5% | 73.7% | 26.3% | Orphaned |
|  | Possums | 5615 | 17.8% | 43.0% | 43.8% | 56.2% | Orphaned |
|  | Small Macropods | 1139 | 3.6% | 8.7% | 47.2% | 52.8% | Orphaned |
|  |  | **13050** | **41.3%** |  | **50.1%** | **49.9%** | **Orphaned** |
| EUTHERIANS | Echidna | 453 | 1.4% | 16.3% | 51.9% | 48.1% | HBC |
|  | Flying Foxes | 2026 | 6.4% | 73.0% | 56.7% | 43.3% | Entanglements |
|  | Microbats | 295 | 0.9% | 10.6% | 65.8% | 34.2% | Cat attack |
|  |  | **2774** | **8.8%** |  | **58.1%** | **41.9%** | **Entanglements** |

|  |  |  |  |  |
| --- | --- | --- | --- | --- |
| *Total / proportion of all admissions* | 31626 |  | 42.6% | 57.4% |
| *Mean proportion of all admissions* |  |  | 46.4% | 53.6% |

<sup>1</sup> Positive outcome includes rehabilitation, sent to carer and released to wild
<sup>2</sup> Mortality includes unassisted death and euthanised on site

### Animal admissions

Mammals represented the majority of admissions to AZWH at 51.1% (*n* = 15,824) (Table 1). Possums (nocturnal marsupials belonging to the Phalangeridae family) were the most commonly admitted multi-species group, with 17.8% (*n* = 5,615) of admissions over the study period. This was closely followed by admissions of koalas (threatened arboreal marsupials; *Phascolarctos cinereus*), at 11.4% (*n* = 3,590), making them the most commonly admitted single species (Table 1, Figure 2a). Eastern grey kangaroos (*Macropus giganteus*) and small macropods (a multi-species group comprising wallabies and pademelons in the Macropodidae family; Supporting File 1) comprised 3.7% (*n* = 1,165) and 3.6% (*n* = 1,139) of all admissions, respectively. Flying foxes (*Pteropus alecto* and *P. poliocephalus*) were the main eutherian mammal admitted (*n* = 2774; 8.8% of admissions), and the fourth most commonly admitted taxa overall (6.4%; *n* = 2,026) (Table 1).

**Figure 2:**
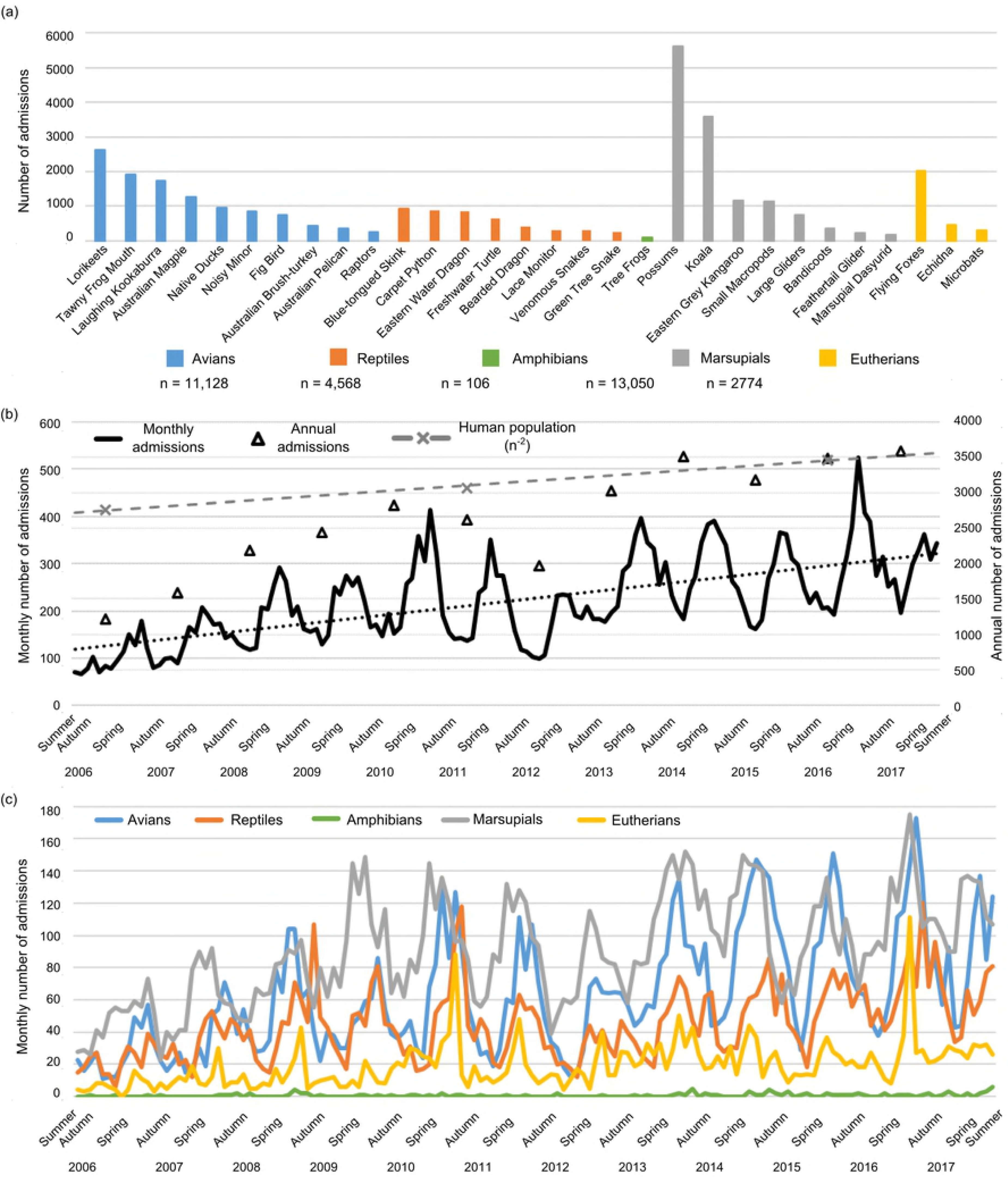
Animal admissions to the Australia Zoo Wildlife Hospital between January 2006 and December 2017 (inclusive). (a) Number of admissions per species or multi-species group. Taxa are ordered within their animal groups by abundance. Taxa are coloured based on higher classifications; see legend. (b) Total admissions per month (left axis) and per year (right axis). The increase in human population in the region is also overlaid (grey dashed lines); one one-hundredth of the total is represented (right axis). (c) Number of monthly admissions per animal group. Taxa are coloured based on higher taxonomic classifications; see legend.

Avians were the second most admitted animal group, accounting for 35.2% (*n* = 11,128) of all admissions (Table 1). The most commonly admitted avian species were lorikeets (colourful psittacines common to Eastern Australia; *n* = 2,625) accounting for 23.6% of avian admissions and 8.3% of admissions overall, and tawny frogmouths (nocturnal birds related to nightjars; *Podargus strigoides; n* = 1,920); whilst high numbers of laughing kookaburras (Dacelo *novaeguineae;*the largest species in the Kingfisher family) and Australian magpies (Gymnorhina *tibicen;* omnivorous passerine songbirds) were also admitted (*n* = 1,741 and *n = 1,263, respectively*).

The reptile group contributed 14.4% (*n* = 4,568) of all admissions, represented by six individual species and two multi-species groups (Table 1, Supporting File 1). Blue-tongued skinks (short legged diurnal lizards; *Tiliqua scincoides*), carpet pythons (large semi-arboreal pythons with a wide distribution; Morelia spilota) and eastern water dragons (arboreal lizards in the Agamidae family; *Intellagama lesueurii)*were the three most commonly admitted reptilian taxa, together comprising 8.5% of all admissions (*n* = 930, 888 and 856, respectively; Table 1, Figure 2a). The remaining 0.3% (*n* = 106) of admissions were attributed to amphibians, represented in our study only by tree frogs (*Litoria caerula* and *Litoria gracilenta*) (Table 1).

We observed a steady increase in the total number of admissions over the study period, with almost a 3-fold increase in annual admissions from 2006 (*n* = 1,216) to 2017 (*n* = 3,582) (Figure 2b, Supporting Table 1, Supporting Figure 1a). The average annual admission rate equated to 2,635.5 animals per year (±744.8). The number of admissions of each animal group also increased steadily, with avians and marsupials showing the greatest increases in admission, at more than 300% throughout the study period (avians; n = 318 to 1,147 and marsupials; n = 562 to 1,505) (Supporting Table 1, Supporting Figure 1, Supporting Figure 2a and 2d).

Seasonal admission trends were apparent in the dataset: the greatest number of admissions occurred annually in spring, with a mean difference of 356.8 (5.5%) from autumn (p < 0.001). Interestingly though, each animal group exhibited a different seasonal profile. Mean bird admissions were highest in spring, as were mammal admissions, while reptile admission peaks occurred largely in summer (Figure 2c, Supporting Figures 1 and 2).

### Causes for admission

Causes for admission (*n* = 31,626) are summarised in Table 2 and Figure 3. The most common CFA was ‘hit by car’ (HBC), accounting for 10,973 admissions (34.7%), followed by ‘orphaned/dependent young’ (24.6%; *n* = 7,771), ‘overt signs of disease’ (9.7%; *n* = 3,057), ‘dog attack’ (9.2%; *n* = 2,913), ‘entanglement’ (7.2%; *n* = 2,274) and ‘cat attack’ (5.3%; *n* = 1,667). These six causes together constituted 90.6% of all admissions (28,655/31,626) and accounted for 64.4% to 100% of admissions for individual taxa. Only four CFA affected all 30 study taxa (abnormal animal location, dog attack, orphaned young, and overt signs of disease), with the remaining CFA applicable for 1 to 29 species or groups (mean 20.5) (Table 2).

**Table 2:**
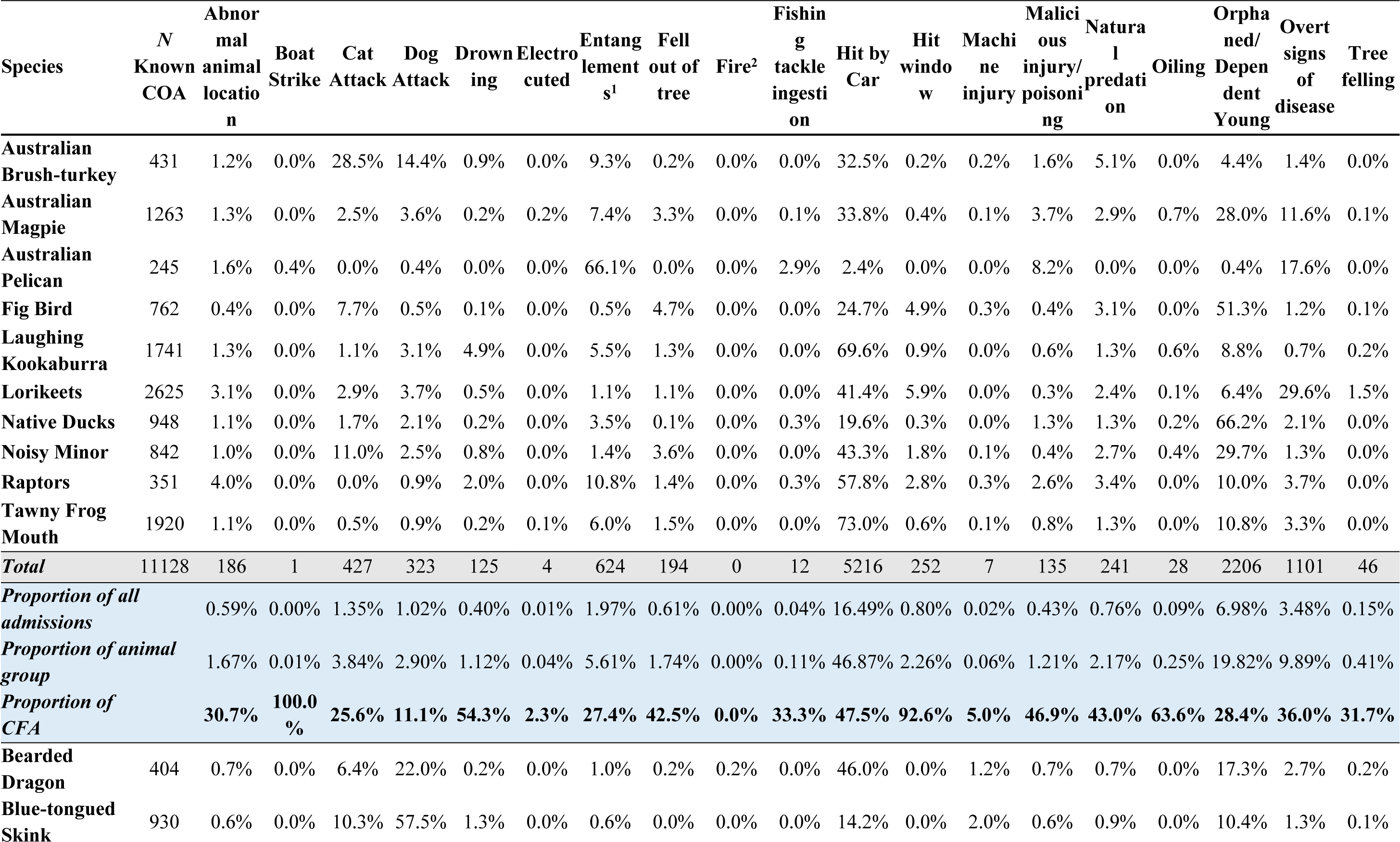
Admissions to AZWH in each CFA, presented as proportion of each species or multi-species group.

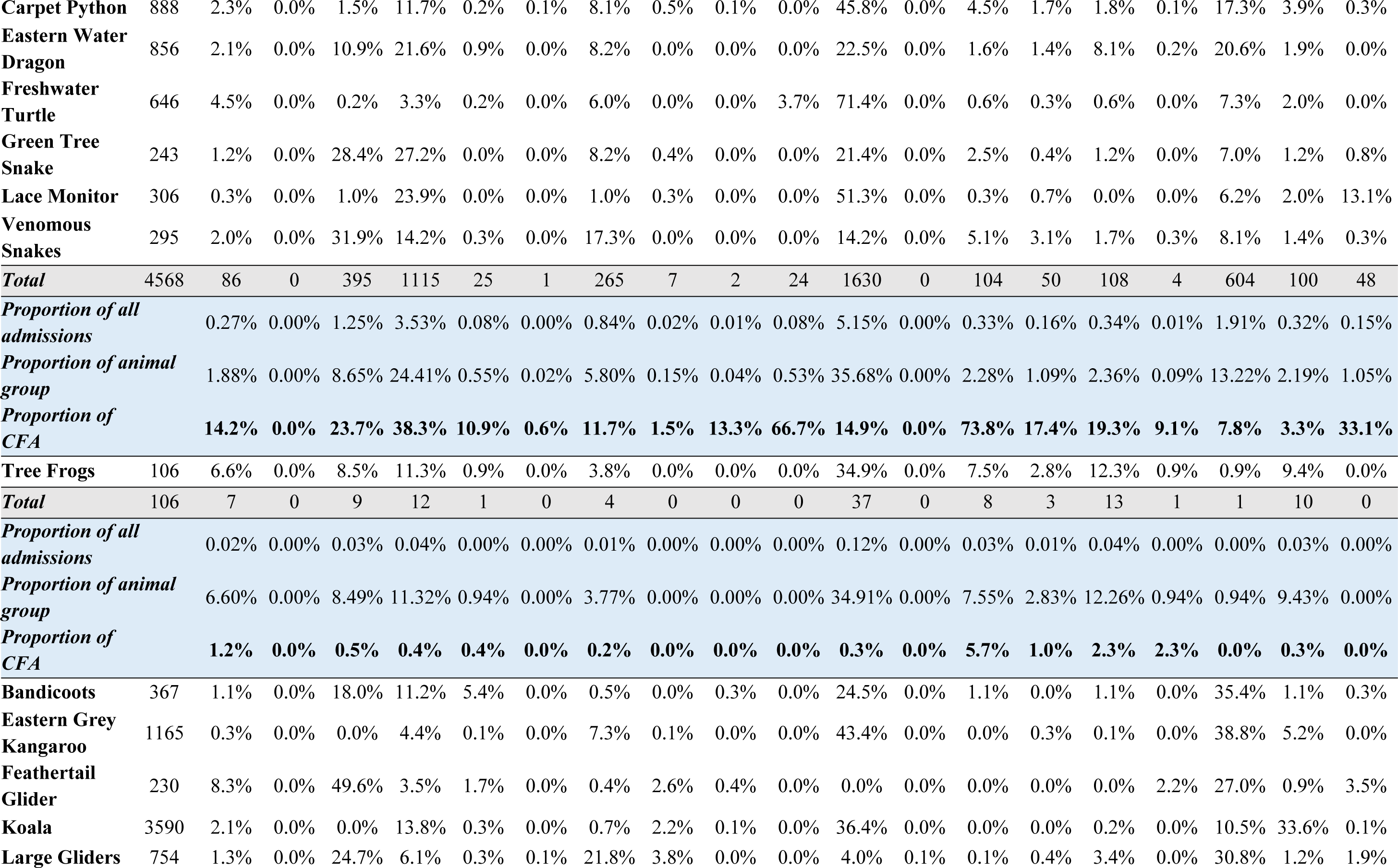

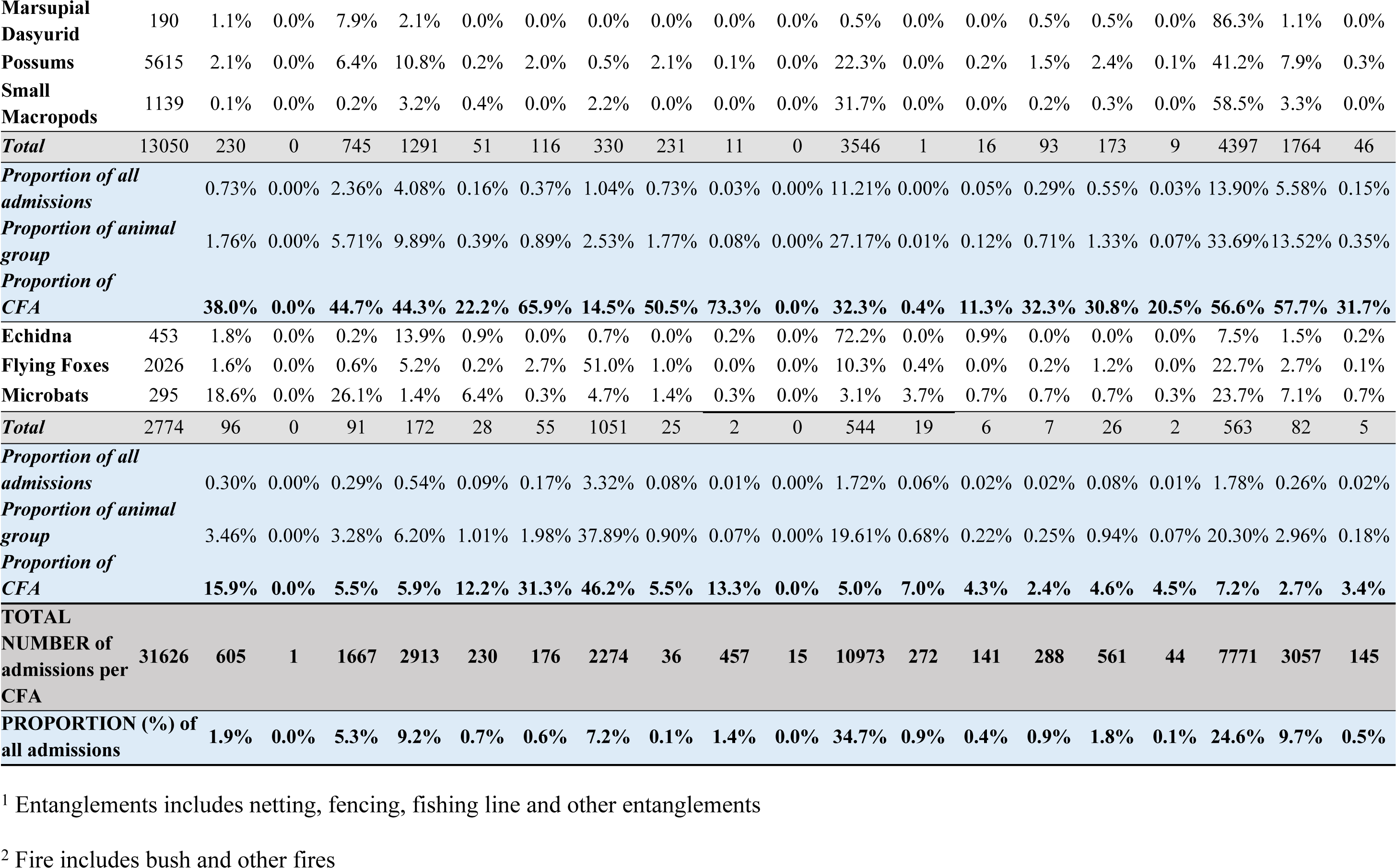

**Figure 3:**
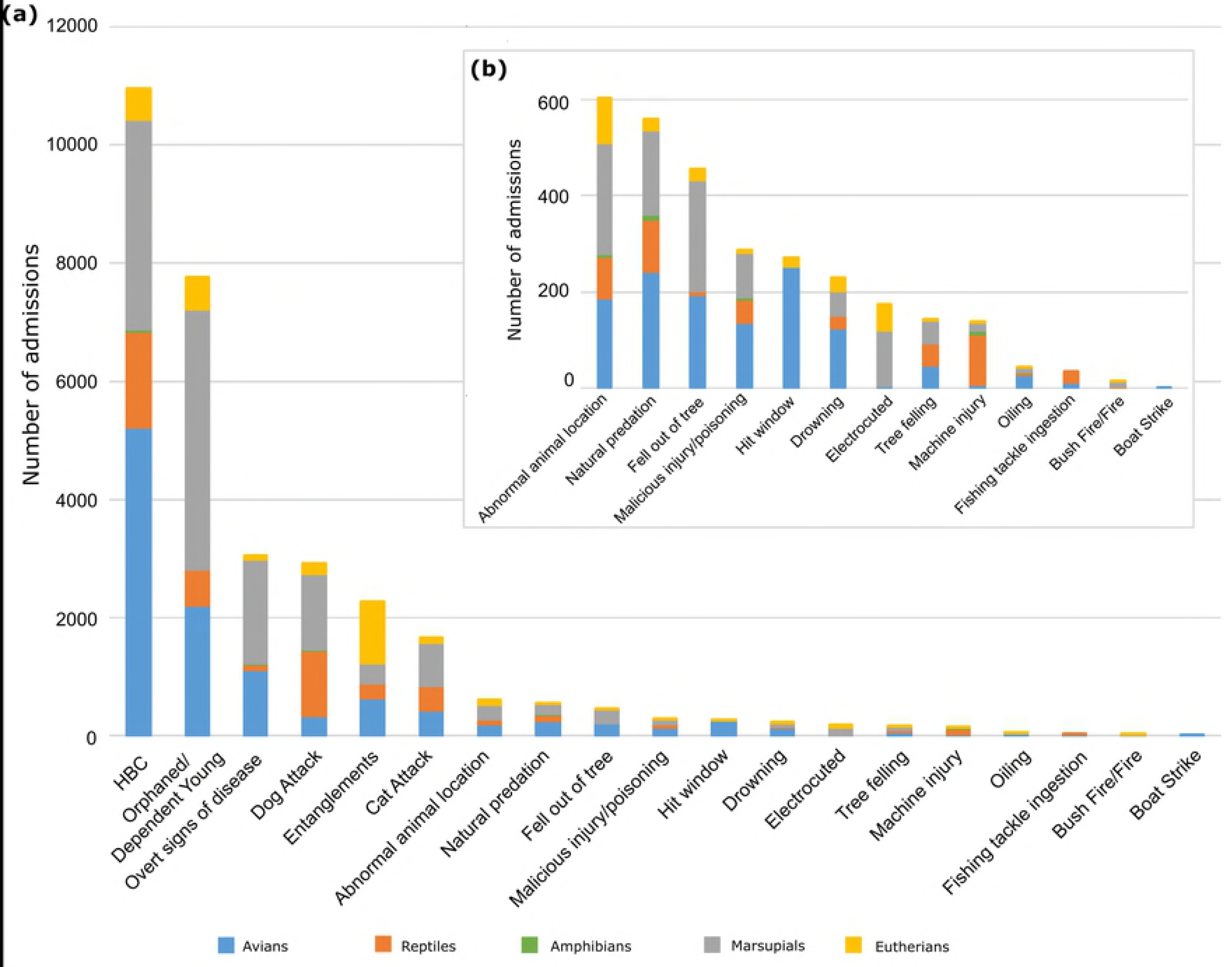
Admissions to AZWH in each CFA. All CFA are represented in descending order on the main graph (a), whilst admissions in categories that are not one of the top six CFA are provided on an additional graph, inset (b). Taxa are coloured based on higher taxonomic classifications; see legend.

Car strikes were the leading cause for admission of 16 out of 19 taxa (Table 1). Avians were the most common group admitted for road trauma (16.5% of all admissions) with 47.5% (5,216/11,128) of avians admitted in this CFA. This mainly comprised tawny frogmouths, laughing kookaburras and lorikeets, which each had over 1,000 admissions (Figure 3, Supporting Table 2). Marsupials and reptiles were also heavily affected by car strikes, accounting for 11.2% and 5.2% of all admissions, respectively (Table 2) with approximately a third of all marsupials (27.2%) and reptiles (35.7%) admitted for this affliction (Figure 3, Table 2). More specifically, koalas and possums together accounted for over 70% of marsupial car strikes (2,558/3,546) whilst freshwater turtles accounted for the highest proportion of reptile car strikes (28.2; 461/1630).

The second highest admission category was ‘orphaned or dependent young’, which accounted for 24.6% of all admissions (*n* = 7,771). Marsupials were most frequently admitted in this category (56.6% of orphaned admissions; n = 4,397; Table 2, Figure 3), with possums alone contributing over half of these (2,314/4,397). Avians together contributed a further 28.4% (*n* = 2,206), mainly consisting of native ducks (*n* = 628).

‘Overt signs of disease’ was one of four CFA shared by all studied species and was the third highest CFA overall (Figure 3). This CFA accounted for high proportions of koala (e.g. chlamydial disease) and lorikeet (e.g. lorikeet paralysis) admissions, at 33.6% (*n* = 1,207) and 29.6% (*n* = 777), respectively. Overt signs of disease also accounted for 17.6% of Australian pelican admissions (e.g. botulism-like symptoms).

‘Dog attack’ was the fourth most common CFA (9.2% of admissions). Marsupials made up the largest proportion of dog attack admissions (44.3%; n = 1,291) and was the CFA for 9.9% of marsupials. Dog attacks accounted for 10.8% to 13.8% of possum, bandicoot and koala admissions (Table 2). Reptiles comprised a further 38.3% of dog attack admissions, with 24.4% (*n* = 1,1115) of reptiles admitted for this reason (Table 2, Supporting Table 2). In particular, 57.2% of blue-tongue skink admissions (*n* = 535) were due to dog attacks.

‘Entanglements’ (e.g. fence or fruit netting entanglements) accounted for 7.2% of all admissions (*n* = 2,274). Eutherian mammals made up 46.2% of all entanglement admissions (Table 2, Figure 3). This mainly consisted of flying foxes (*n* = 1,034), for which entanglement accounted for 51.0% of admissions. Avians comprised a further 27.4% of entanglements (*n* = 1,253), with a heavy proportion of Australian Pelicans admitted following entanglement (66.1%; *n* = 162; Table 2, Supporting Table 2). Entanglements also represented a sizeable proportion of large glider admissions (21.8%; *n* = 164). This multi-species group consists of the greater glider, squirrel glider and sugar glider, which are comparable in size to flying foxes.

‘Cat attack’ rounded out the top six CFA at 5.3% of all admissions (*n* = 1,667). Cat attacks accounted for 49.6% of feathertail glider admissions (*n* = 114), and over 20% of admissions of Australian brush turkeys, green tree snakes, venomous snakes, large gliders and microbats (Table 2). Over 8% of both reptiles and amphibians were admitted due to cat attacks (Supporting Table 2).

Some animals had unique or specific CFA that were distinct from the top six CFA. Reptiles were commonly admitted for ‘machine injury’, which includes incidents involving lawn mowers, grass cutters, whipper snippers, chainsaws, tractor slashers etc (Figure 3). Carpet pythons were also highly represented (*n* = 40) in this category (Table 2). The most common CFA for amphibians was HBC (*n* = 37), but they were also prone to dog attacks and ‘natural predation’ (native predator attack resulting in injury; *n* = 12 and 13, respectively). In fact, natural predation accounted for 12.3% (*n* = 13) of amphibian and 8.1% (*n* = 69) of eastern water dragon admissions (Table 2, Supporting Table 2). Lace monitor admissions were primarily the result of tree-felling (13.1%; *n* = 40), which results in injury or displacement. ‘Abnormal animal location’ was a common CFA for microbats and feathertail gliders (18.6%; *n* = 55 and 8.3%; n = 19, respectively; Table 2), whereby they may be found on the ground, in unsuitable locations within building infrastructure, or other locations compromising their welfare. A small percentage of animals (0.9%, 288/31,626) were admitted for ‘malicious injury or poisoning’, where injury or illness was suspected due to a malicious act. Australian pelicans appeared to be overrepresented in this category (*n* = 20, 8.2% of pelican admissions), though this may be the result of assignment of some pelicans affected by botulism-related disease to this category (Table 2). Eight animal species were affected by ‘fire’, which includes bush fire and other fire, and ‘electrocution’. Admissions resulting from fire-related events were mostly restricted to mammals (Table 2). Electrocution largely affected arboreal animals from all groups, with possums and flying foxes most commonly admitted under this category. Birds and bats were infrequently admitted for hitting a window, whilst ‘fishing tackle ingestion’ admissions were restricted to birds and freshwater turtles (Table 2). Laughing kookaburras were the most commonly admitted species in the ‘drowning’ (*n* = 86) and ‘oiling’ categories (*n* = 11) and lorikeets in the ‘hit window’ (*n* = 155) category.

Consistent with the overall increase in admissions over time, admissions due to each of the top six CFA increased considerably over the study period, with these six CFA increasing by up to ten-fold between 2006 and 2017 (Figure 4a, Supporting Figure 3).

**Figure 4:**
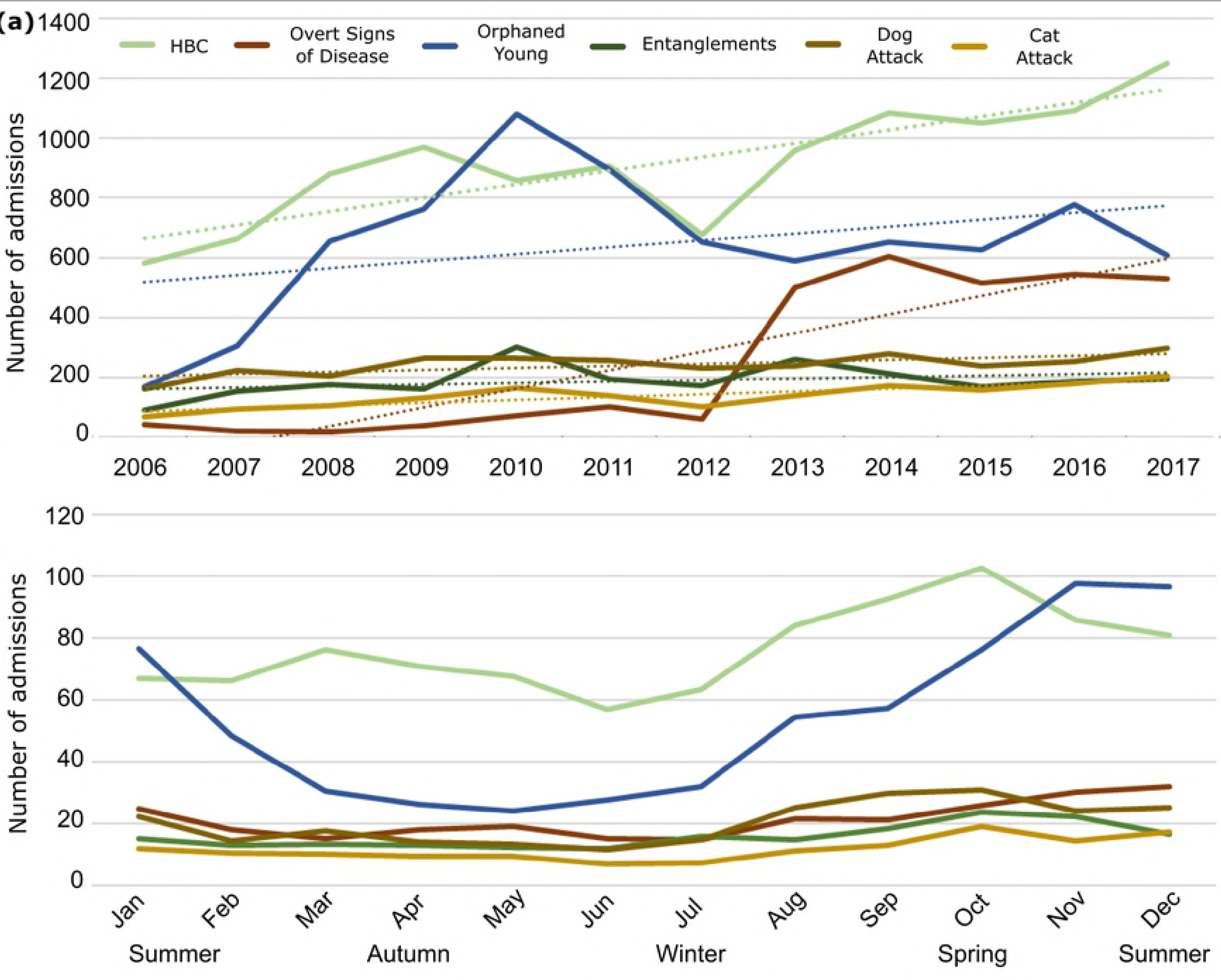
Annual (a) and seasonal (b) animal admissions to AZWH for the top six CFA. Trend lines are included in (a) to highlight the overall increase in admissions over the study period. See legend for CFA categories.

Some CFA exhibited cyclic trends (Figure 4b, Supporting Figure 4). Admissions of orphaned animals were clearly seasonal (admissions in spring were statistically different from admissions in autumn, winter and summer (*p* = 0.001, 0.018, 0.016, respectively; Figure 4b), with avian and marsupial admissions increasing from late winter, and remaining high throughout spring and summer (Supporting Figure 4). Entanglements peaked in spring, and dog attack admissions were highest overall during late winter and spring.

### Outcomes of admission

Animal outcomes following admission were grouped simply into either ‘positive ‘outcome’ or ‘mortality’. Positive outcomes included release into wild or into care, whilst mortality encompassed both natural death and euthanasia on welfare grounds.

Mortality was listed as the outcome for the majority of animals (57.4%; *n* = 18,153), with an average mortality rate of 53.6% (Table 1, Figure 5). Overall mortality among birds and reptiles was slightly greater than the average (55.7% and 57.4%, respectively; Table 1), whilst mortality in amphibians was highest at 67.9% (72/106). Lorikeets had the highest mortality rate at 75.2%, whilst Australian pelicans had the lowest mortality rate at 16.7% (Figure 5a).

**Figure 5:**
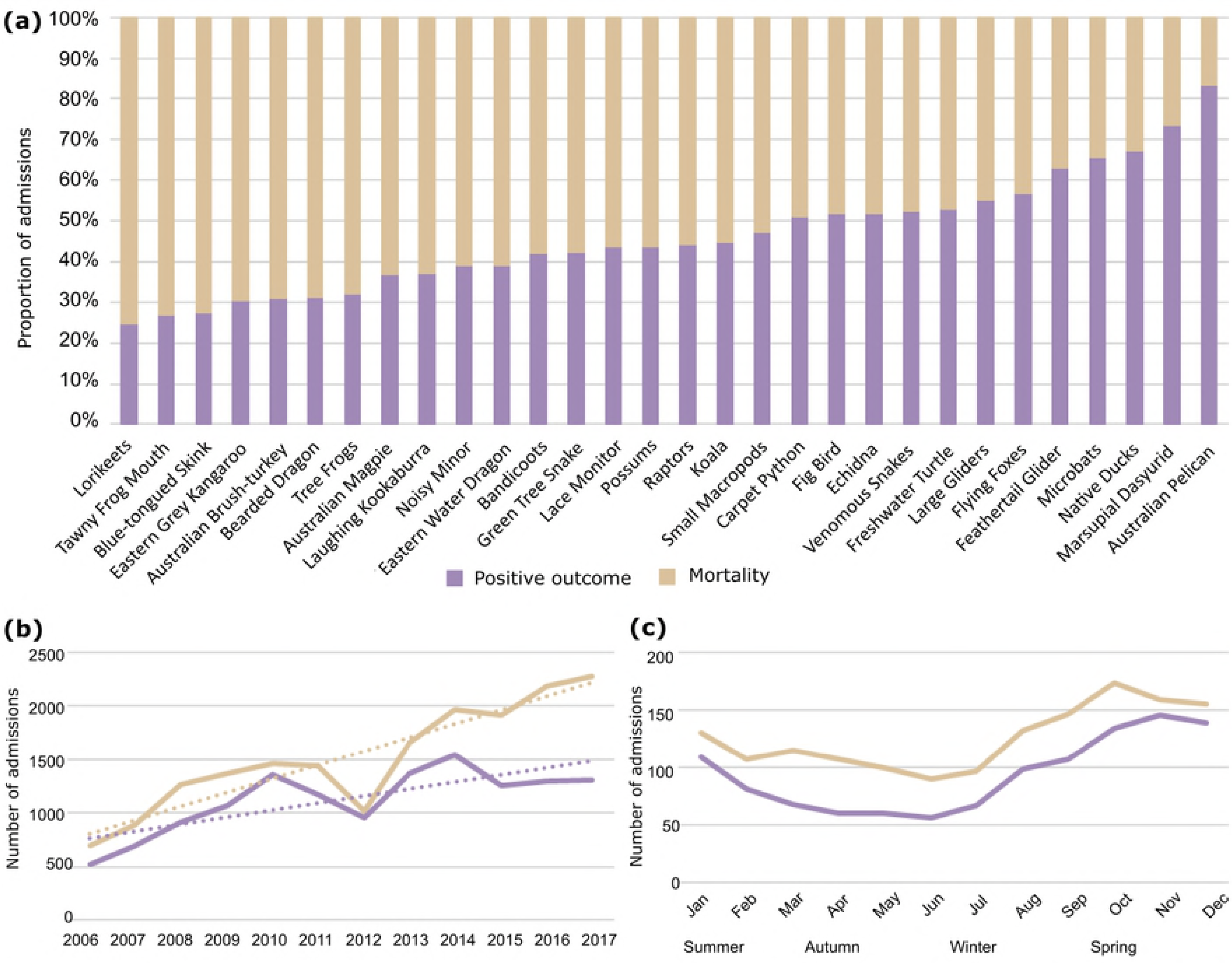
Outcomes of admission to AZWH between 2006 and 2017. (a) Proportion of total admissions for each species or multi-species group. Total annual (b) and mean monthly/seasonal (c) admissions resulting in positive outcomes and mortality. Trend lines are included in (b) to emphasise the increasing disparity between positive outcome and mortality over time.

Deaths due to HBC accounted for 26.0% of all admissions (8,208/31,626; Table 3). Mortality rates among individual species attributed to HBC ranged from 44.4% (microbats; 4/9) to 92.5% (eastern grey kangaroo; 468/506), with an overall mortality rate of 74.8%, and a mean mortality rate of 69.1% (Table 3, Supporting Figure 5). HBC also had the highest odds ratio for mortality at 3.3 (Table 4).

Dog attacks had the highest mean mortality rate at 72.7%, with 80.8% and 80.4% mortality rates in avians and reptiles, respectively (Table 3). The relative risk of dog attack was second only to HBC, at 1.333, and the odds ratio for mortality ranged from 0.542 in amphibians to 3.741 in reptiles (Table 4, Supporting Table 4). Cat attacks also resulted in high mortality rates, ranging from 39.1% in green tree snakes to 81.3% in native ducks (with the omission of animals that had fewer than 4 cat attack admissions; Table 3, Supporting Figure 5). The relative risk for cat attacks (1.126) was lower than that for dog attacks (Table 4).

**Table 3:**
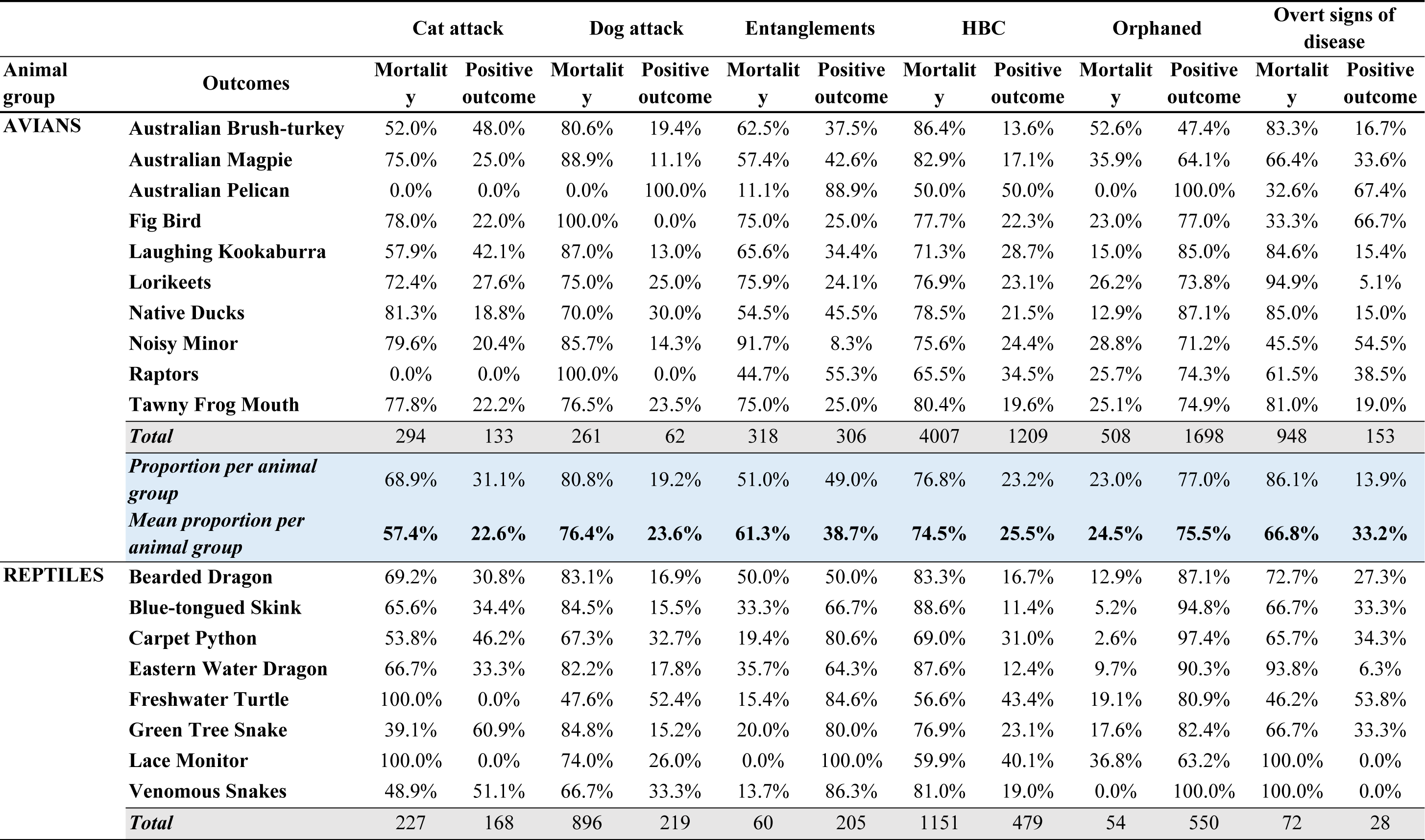
Outcomes of the top six CFA for each species or multi-species group.

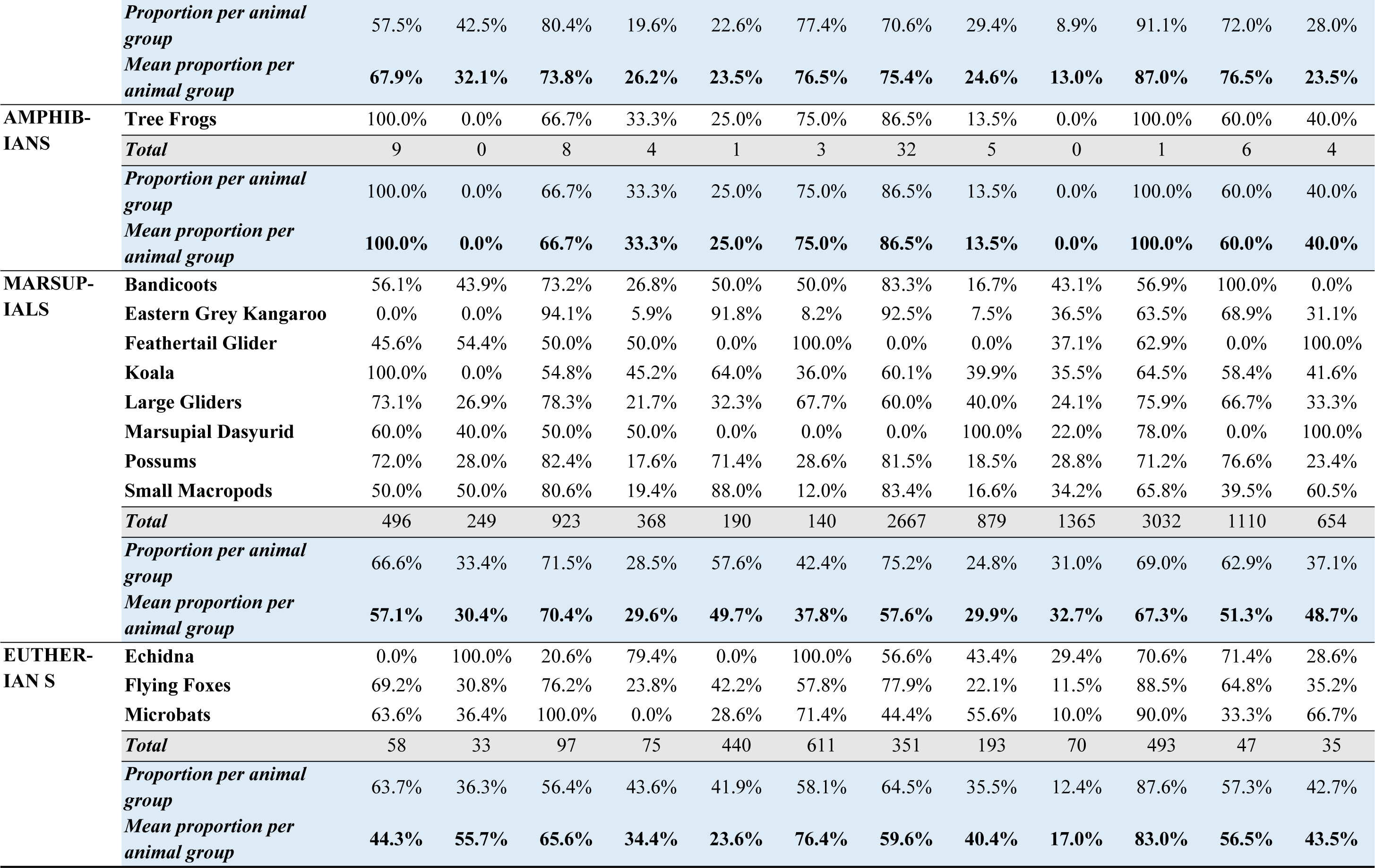

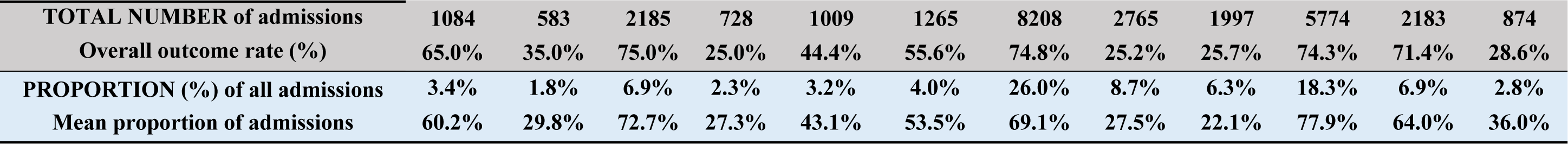

**Table 4:**
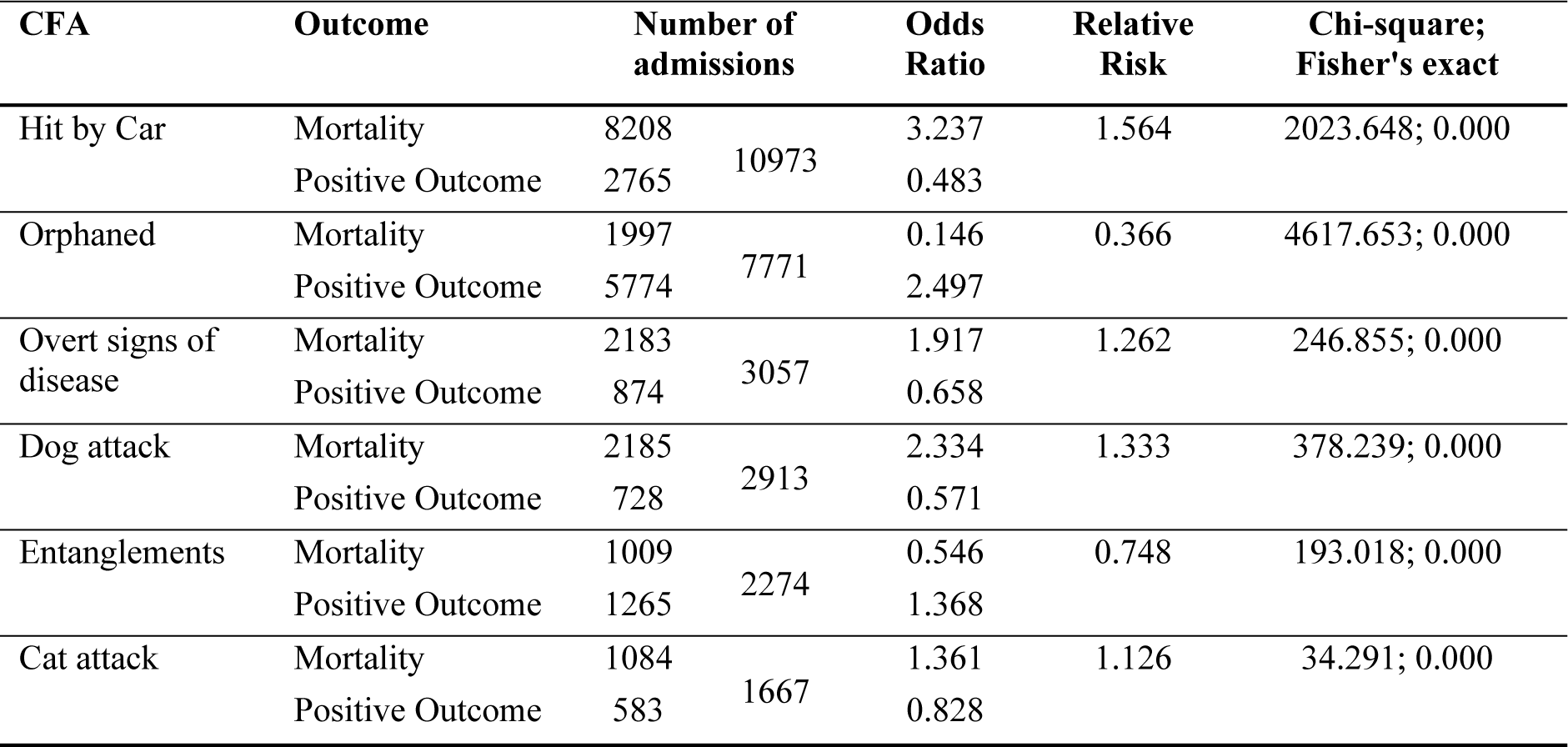
Odds ratio and relative risk analysis for the top six CFA.

| CFA | Outcome | Number of admissions |  | Odds Ratio | Relative Risk | Chi-square; Fisher's exact |
| --- | --- | --- | --- | --- | --- | --- |
| Hit by Car | Mortality | 8208 | 10973 | 3.237 | 1.564 | 2023.648; 0.000 |
|  | Positive Outcome | 2765 |  | 0.483 |  |  |
| Orphaned | Mortality | 1997 | 7771 | 0.146 | 0.366 | 4617.653; 0.000 |
|  | Positive Outcome | 5774 |  | 2.497 |  |  |
| Overt signs of disease | Mortality | 2183 | 3057 | 1.917 | 1.262 | 246.855; 0.000 |
|  | Positive Outcome | 874 |  | 0.658 |  |  |
| Dog attack | Mortality | 2185 | 2913 | 2.334 | 1.333 | 378.239; 0.000 |
|  | Positive Outcome | 728 |  | 0.571 |  |  |
| Entanglements | Mortality | 1009 | 2274 | 0.546 | 0.748 | 193.018; 0.000 |
|  | Positive Outcome | 1265 |  | 1.368 |  |  |
| Cat attack | Mortality | 1084 | 1667 | 1.361 | 1.126 | 34.291; 0.000 |
|  | Positive Outcome | 583 |  | 0.828 |  |  |

The overall rate of positive outcomes was 42.6% (*n* = 13,473), and the average rate of positive outcomes ranged from 32.1% for amphibians to 58.1% for eutherian mammals (Table 1, Table 3, Supporting Table 3). Marsupials had 50.1% positive outcomes and 49.9% mortality across all CFA (Table 1).

Orphaned or dependent young carried the highest rate of positive outcomes (77.9%), which was high in all groups, ranging upwards from 69.0% of marsupials, and the associated relative risk of mortality for all species was only 0.366 (Table 3, Table 4). The relative risk of mortality was lower than average in avians, reptiles and eutherians (Supporting Table 4). Entanglements had a relatively high positive outcome rate, at 53.5% on average, with reptiles and eutherians exhibiting very high positive outcome rates (76.5% and 76.4%, respectively) (Table 3, Supporting Table 3). Relative risk of mortality was also low at 0.748, although the risk was higher for marsupials and eutherians (Table 4, Supporting Table 4).

Overall, increases in annual admissions were mirrored by increases in mortality rate (Figure 5b), however, this was not accompanied by a change to the average annual mortality rate. There were no prominent seasonal differences between positive and negative outcomes overall (Figure 5c).

## Discussion

Native wildlife faces an ever-increasing range and magnitude of threats with the continuing increase of human population, associated urbanisation and anthropogenic-driven climate change being of immediate concern. Several studies have characterised declines in particular species or animal groups, whilst others have examined the impacts of a specific threat in a single biogeographical location, yet few have quantified the factors contributing to morbidities and mortalities longitudinally across a wide taxonomic range of native fauna.

This study has the widest breadth of any longitudinal analysis to date on the animals admitted to a WRC. It examines and critically analyses trends in admissions, causes for admission and animal outcomes over a twelve-year period at a WRC in South-East QLD, Australia. We observed a mean annual admission rate of 2,635 animals for the dataset examined, comparable to some previous studies in Europe, Africa and USA [19, 24, 26, 61, 62]. Differences in admission rates between WRCs in different countries or biogeographical areas are largely a consequence of variations in species richness, human population density, local natural and anthropogenic threats, admission capacity and cultural attitudes to wildlife.

### WRC databases provide an opportunity for wildlife monitoring

Mammalian and avian taxa were the most commonly admitted groups in our study, reflecting the abundance and diversity of these groups in South-East QLD. Mammals comprised over 50% (*n* = 15,826) of our dataset, providing a wealth of knowledge regarding the diversity and abundance of these native animals in South-East QLD. Of these, koalas, possums and flying foxes were among the five most admitted animals overall, highlighting the need for us to understand the human-induced pressures placed on these animals. A further 35.2% of our studied admissions were avians. This is considerably lower than other studies that report up to 57.1% [47] in the UK, and even 90% [63] in South Africa, whilst higher than a study from the USA (12.2%) [24].

We expect that these discrepancies are largely due to differences in species richness in SEQ compared to other regions [13, 23, 43, 49, 64, 65]. These differences will inform and influence monitoring efforts and conservation priorities. We focussed only on terrestrial and freshwater species (including avians), omitting marine species as we consider these to be threatened by distinct factors warranting their own analysis.

An overall increase in admissions was witnessed over the study period, which we believe is largely attributed to human population increases, as evidenced by the increase over time of admissions due to human-associated CFA. This is supported by human population growth in the Sunshine Coast region from 236,654 residents in 2006 to 303,389 in 2016 [66]. The population is expected to reach 500,000 by 2031 [67], which we anticipate will result in further increases in wildlife admissions to AZWH.

### Human activities are contributing to the decline of Australian icons

Given their iconic nature as representatives of the unique fauna found in Australia, and “vulnerable” status (up from “least concern” in 2008) [68], the health, welfare and conservation status of koalas continue to be of prime interest for the Australian public and the international community. Koala populations have suffered massive decline over the last 30 years, particularly in QLD, with recent localised population collapses documented [4, 5, 69]. Emphasising the precarious nature of the koala’s survival in South-East QLD, koala admissions were high and constant throughout the study period, consistent with reports from other WRCs [1, 3-5, 69].

Major threats to the koala include habitat fragmentation, road trauma and disease [4, 69]. Land clearing, to facilitate urban expansion and agriculture is also having devastating effects on the welfare of native fauna worldwide [37]. Whilst we did not directly measure habitat fragmentation in our study, most koala admissions were from urbanised areas with high numbers of car strikes, dog attacks and animals found in abnormal locations (e.g. telegraph poles and bridges), demonstrating a clear link between urban encroachment and its effect on koalas. Chlamydial disease is highly prevalent in koalas from South-East QLD and has been identified as a key threat to koala populations [70, 71]. As such, identifying and quantifying the prevalence of chlamydial disease in koalas is vital for ongoing management. Urogenital disease caused by *Chlamydia pecorum* can diminish the fecundity of the population as it can lead to infertility, whilst ocular disease can lead to blindness and increased risk of morbidity. Overt chlamydial disease, in the form of a stained rump and inflamed exudative eyes, is one of the most common reasons for koala rescue and admissions to WRCs in South-East QLD, yet hospital databases may not always accurately capture this as a primary CFA. In 2013, AZWH revised their animal accession/admission data capture and on-site database to enhance both the quality of animal admission data and ability to report on CFA. This process included revisions to CFA categories and the inclusion of a category for ‘overt signs of disease’. As a result, admissions for overt signs of disease appeared to increase markedly from mid-2013 (Figure 4), yet realistically, the prevalence of overt chlamydial disease in koalas was similar to previous years. Our study was able to demonstrate how advances in the accuracy of data recording can result in an improved understanding of true threats to wildlife.

The most commonly admitted multi-species group in this study was possums, which are prolific in South-East QLD and thrive in urban areas. Due to their widespread nature and high density within urban and peri-urban regions, possums are predisposed to anthropogenic-related threats as demonstrated by high numbers of cat attacks, dog attacks and car strikes in this study, all of which resulted in high proportions of mortality (72.0%, 82.4% and 81.5%, respectively).

Another iconic Australian marsupial is the kangaroo. A recent study of eastern grey kangaroo with an overlapping study area, but also encompassing other regions of Australia, found that 42% of studied populations were in decline, with the most prominent impacts found in areas of high, ongoing urbanisation and transport infrastructure development [72]. In support of these findings, within our study area, 43.4% of eastern grey kangaroos were admitted due to car strike, with 92.5% of those incidents resulting in mortality; eastern grey kangaroos had the fourth highest total mortality rate. Interestingly, small macropods fared better than eastern grey kangaroos following car strikes for which they were commonly admitted, with more than double the positive outcome rate (16.6%), which was contrary to expectations as they have similar physiological and behavioural traits other than body size. Overall these results reiterate the substantive negative impacts of building roads through remaining habitat or habitat linkage pathways of animals that are already vulnerable due to previous habitat modification and destruction at a landscape scale, in the absence of the implementation of adequate conservation strategies to mitigate the negative impact. By addressing factors such as vehicle density, vehicle speed, signage, road side habitat and lighting (including daylight savings time) and appropriately designed wildlife corridors, the impact of vehicle collisions can be reduced [3, 16, 73]. However, the rate of human population expansion and urbanisation in our study area, as well as across many global regions, mean that vehicle associated wildlife mortality will still occur and likely constitutes one of the most prominent threats to the persistence of viable wild populations of many taxa.

The highest mortality rates of any taxa in this study were for lorikeets. Rainbow lorikeets are one of the most commonly observed birds in Australia with a natural distribution along the east coast [74] but are actually considered pests in other parts of Australia and New Zealand [75]. Whilst they were most commonly admitted in the hit window category, tree-felling and disease were also common reasons for lorikeets to be admitted. Disease resulted in a 94.9% mortality rate in lorikeets. Two diseases are primarily responsible for this: Psittacine beak and feather disease, a skin disease caused by *Circovirus* that is often fatal [76]; and necrotising enteritis, a gastrointestinal disease caused by *Clostridia spp*[77]. The latter is associated with altered dietary regimes associated with human habitat modification, or in some instances ingestion of inappropriate food directly sourced from humans in the form of garden bird-feeders and human food [77, 78], providing yet another example of the preventable impact of human activities on wildlife.

### Human-related CFA contribute to higher wildlife mortality rates

Unfavourable outcomes were statistically more likely if the CFA was domestic cat or dog attack, car strike or entanglements. The combined average mortality rate of these four human-related CFA was 61.3%., with the relative risk of mortality ranging from 1.3 to 1.6 compared to 0.4 and 0.7 for orphaning or overt signs of disease, respectively. These differences are due to the severity of the trauma caused by cats, dogs, cars and fencing and netting, which reduce the likelihood of successful rehabilitation, and are also likely underrepresented in our data given that orphaning would be in many instances a result of human linked impacts on the parents of orphaned individuals.

Entanglements were one of the human-related CFA responsible for a high proportion of admissions and mortality, again driving home the significant impact human activities have on a diverse range of wildlife. In the case of flying foxes, which were the fourth most commonly admitted taxa in our study, 51% were admitted on the basis of entanglements, which resulted in a 57.6% mortality rate. Whilst we grouped all types of entanglements due to insufficient data resolution within the source database, a recent study in Victoria showed that a high proportion of animals were admitted due to fruit netting entanglements (36.8%), where up to 56.1% of each entanglement subcategory resulted in mortality [6]. This was one of the highest mortality rates in our study and suggests that changes in land management practices may be the most effective way of ameliorating native wildlife mortality associated with entanglement, particularly for terrestrial taxa. Within the study region, several local councils have initiatives such as ‘land for wildlife’, partly aimed at converting conventional barbed-wire livestock fencing into wildlife friendly options, as well as reducing the use of monofilament netting which present entanglement risk to taxa such as flying foxes [79]. However, these efforts rely on goodwill from landholders, and there is no legislative requirement at either the local, state or federal level to enforce such practices. This highlights the need for consistent, overarching policy to guide land management practices toward mitigation of unnecessary risk to native fauna.

Estimates in the USA place annual cat-related predation in the billions [21, 22], and predation of native animals by both feral and domestic cats in Australia is similarly devastating. For example, predation by feral cats has resulted in the early localised extinction of indigenous wildlife such as the western quoll (*Dasyurus geoffroii*) and golden bandicoot (*Isoodon auratus),* from islands off Western Australia [80], with more recent declines in numbers of other marsupials such as the northern brown bandicoot (*Isoodon macrourus*) in Northern Australia [9, 81, among other examples [81, 82, 83]. Cats are ubiquitous in Australia, with millions kept as pets that are permitted outdoors, and others free-ranging in urban environments and the wild [82-84]. Cat attacks have particularly serious effects on birds and reptiles, and microbats are also especially susceptible to cat trauma, demonstrated by 63.6% mortality in our study, 28.7% of bat casualties in a study in Italy [85] and around half of the traumatic deaths of bats found in Germany [86]. The cat attack admissions figures in our dataset are deceiving as we omitted animals that were DOA. Cats are generalist predators that are known to consume prey, which has also been documented in Northern Australia: birds, small mammals and small reptiles are common food sources when available [87]. Such mortalities were not captured in our dataset. Further, cat removal measures have resulted in reversal of population declines in some areas [88, 89] suggesting that such measures may be successful elsewhere. The culling of dingoes in many Australian jurisdictions has also been demonstrated to be detrimental to ecosystem functioning, as they act as top predators, often minimising the negative effects of feral mesopredators such as cats and foxes [90-93]. Further, cats do not only prey on native fauna but may also out-compete smaller bodied native predators such as quolls for resources [89], proving another, indirect, effect of the negative impact of such introduced species on our native fauna.

Dog attacks were another CFA resulting in significant mortality, with reptiles highly represented in this category. This is in agreement with another Australian study that showed around 49.2-52.4% of admissions of bluetongue lizards, which are common in backyards, were admitted following dog attacks, and 70% of all dog attack admissions did not recover [8]. A study in Tennessee however, reported far fewer admissions (only 6.1%) of reptiles due to dog attacks, where “human-induced trauma” was listed as the most common CFA for reptiles [24]. Dog attacks were also responsible for high mortality rates of koalas in our study. This is another example of the value of local wildlife monitoring to ascertain the specific threats faced by wildlife in distinct regions.

### The influence of animal morphology and behavioural traits on predisposition to threats

Habitat characteristics, foraging practises, circadian movement patterns, size and other behavioural traits appear to predispose some taxa to certain threats, which are augmented by human-induced habitat alteration in the absence of suitable measures for impact reduction. The CFA for which this appears most clear is HBC, which was the leading CFA in our study. A detailed review of road trauma throughout Europe reported on average 2 to 8.5 million road kills per year among birds, reptiles and mammals (particularly ungulates) in countries such as the Netherlands, Belgium and Sweden [15]. The authors suggested that these animals are predisposed to vehicle collisions due to behavioural and ecological factors. A recent review of the propensity of wildlife to suffer from car strikes highlighted the increased risk for omnivorous avian taxa [94], which can be correlated in our study with the high rate of car strikes for tawny frogmouths, which are nocturnal omnivores with a tendency to hunt for insects that are attracted to car headlights on the road. Other avian species utilise roadside telegraph poles and fences as vantage points for hunting, further predisposing them to vehicle strikes [58].

Similarly, hedgehogs, which are nocturnal animals with morphological and behavioural traits resembling echidnas, (i.e. relatively slow movement, poor eyesight, limited defence against car strikes) have been documented to be profoundly affected by car strikes in the UK, with admission and mortality rates due to car strikes of 10.3% and over 85%, respectively [50]. In our study echidnas had a much higher admission rate due to car strikes (72.2%) with a corresponding mortality rate of 56.6%. Other taxa, such as herpetefauna are predisposed to car hits as they may be drawn to the microclimate of a warm road, or they may be migrating to or from a hibernation site [51]. Turtles are also disadvantaged at evading car strikes due their slow speed, as evidenced by previous Australian research, that reported an 82.3% admission rate of Long-necked turtles over a 13-year study, with an overall mortality rate of 60.9% after impact with a motor vehicle. This is comparable to the mortality rate of freshwater turtles in our study at 56.6%, as well as the morbidity/mortality rate reported for three turtle species at a WRC in Virginia [95]. These findings are also consistent with a study that showed that maximum sprint speed may be a determinant of an animal’s ability to evade injury or mortality associated with car strikes [94]. Further, Heigl *et al* reported a higher number of road-killed amphibians and reptiles on agricultural roads than municipal roads. Whilst we didn’t measure this in our study, our common admissions area does include rural and bushland zones, so a similar trend may be apparent in our study.

We saw prominent differences in the admission and outcome rates of predatory, aggressive, or territorial birds versus more placid birds. For example, there were only 351 admissions of raptors, which is a grouping of 17 species. Raptors were the only birds apart from pelicans and noisy minors that were almost never admitted due to cat or dog attack, with low admissions most likely to their low relative abundance, coupled with their behavioural characteristics, which comprise ambush attack on prey from high vantage points, with little time spent in vulnerable positions. Conversely, noisy minors, although smaller in body size than raptors, are gregarious and territorial, forming colonies that can contain hundreds of birds providing a means of communal territory defence, which could explain the relative paucity of dog and cat attacks. These behavioural traits may also influence people’s perceptions of the value of certain wildlife and the likelihood of presenting them to a WRC, for example in the case of noisy minors.

### Severe weather events result in spikes in admissions

Besides an overall increase in admissions over the course of our study, we observed several distinct peaks in total admissions (2010, 2014, 2016, 2017) that may be correlated with severe local weather events affecting the region of South-East QLD, Australia. December 2010 recorded the “wettest December on record” with widespread heavy rainfall and thunderstorms, culminating in one of the most significant flood events in QLD’s recorded history [96]. Flood events damage animal habitat and alter animal movement and behavioural patterns, often resulting in mortality, displacement, injury, stress or disease. We observed an expected increase in orphaned cases in December 2010, particularly for birds and marsupials. Reptile admissions did not show the same trend, which may reflect the ability of snakes in particular to traverse floodwaters by swimming. Animals capable of climbing, which are heavily represented in our dataset by arboreal marsupials, may not have been as heavily affected by flooding, but thunderstorms, such as the ‘super-cell’ that affected the city of Brisbane in South-East QLD (Figure 1) in November 2016 [97], likely resulted in mass animal displacement and injury, evidenced by a similar increase in orphan cases at that time. The same month also saw a heatwave in Kilcoy (~40 km west of AZWH), which, combined with recent land-clearing in the area, resulted in mass morbidity and mortalities of flying foxes.

Unusually dry and hot months were seen in Spring 2014, with QLD temperature records broken through October and November 2014 following ongoing and widespread drought [97], prior to a damaging super cell storm in Brisbane at the end of November with heavy wind gusts and large hail stones [99]. These events coincide with peaks in avian and marsupial admissions. Similarly, 2017 was Australia’s third-warmest year on record, with persistently warmer than average days year-round [100]. High ambient temperatures cause morbidity and mortality due to heat stress, whilst prolonged drought destroys habitat and limits food and water sources. Alongside the more obvious and conspicuous threats associated with human activities, such as car strikes, these results highlight that anthropogenically induced climate change will likely exacerbate threats to wildlife, due to the predicted higher frequency of severe weather events that have not been as prevalent in the recent evolutionary history of Australian fauna.

### Seasonality of admissions

Previous studies have shown that admissions to WRCs are markedly higher throughout the breeding season of included taxa (commonly occurring in spring) [5, 13, 47, 85, 101, 102]. As the weather begins to warm, many native species begin courtship and mating, prior to nesting, giving birth and carrying young. Some young may also go through weaning, and later disperse during the spring and summer months. Studies of birds and mammals in WRCs in South Africa and Colorado exhibited peaks in overall and orphaned/juvenile admissions during their common breeding season [17, 63]. The same trend was also apparent in a 15-year longitudinal study of little owls in Spain in which orphaned young were the most common CFA overall [13]. Furthermore, peak admissions were also reported for reptiles in late spring in Victoria, Australia [8]. We observed similar increases in admissions in our study, with higher admission rates overall during the spring months (September, October, November; mean difference of 356.8 from autumn; p < 0.001). The precise timing of species-specific admission peaks varied between animal groups which is likely a reflection of the relative length of breeding seasons, mating and nesting habits, gestation period, and time to independence for different taxa. Peak periods of juvenile dispersal also coincide with influxes of holidaying families and tourist drawn to the Sunshine Coast region, for the summer Christmas holiday period (December and January), resulting in increased human activity and motor vehicle use. We believe this cyclical, transient population increase and its effects on wildlife can be used to predict the long-term effects of ongoing urbanisation in the area and further highlight the need for proactive conservation management to be a paramount consideration in short and long term town planning for the region.

### Limitations and future directions

The primary but unavoidable limitation of this study lies in the fact that causes for morbidity that occur in close proximity to or are directly due to human activities are strongly selected for in our study. Car strikes, entanglements, domestic dog and cat attacks, window hits and mower strikes are all examples of this bias, with displacements from normal habitat also potentially bringing animals into closer proximity with humans and their activities. Further, charismatic and non-threatening animals such as possums and several birds are more likely to be admitted to WRC’s than seemingly dangerous, unpredictable or large animals such as snakes, kangaroos and large reptiles. These limitations are common among these types of studies and have been raised by other authors [19]. Importantly, they highlight the significant impact of human activities on wildlife welfare and the need for awareness and education. There may also be a related bias toward diurnal animals, as humans are more likely to present injured animals during the day.

Some CFA categories are likely under-represented or may be mis-categorised. One example is cat attack admissions, whereby the devastating effects of domestic and feral cat predation on Australian wildlife are well established [81], however their mode of predation often results in mortality or injury in a manner that does not result in WRC admission [84], or were omitted entirely from our study because they were DOA. This is also likely to be true of fox predation, leading to under-representation within this dataset. Disease may also be under-represented: for example, reptile viral disease is often undetected if funding is unavailable to carry out specific diagnostic tests, and botulism in birds may be placed into the poison category.

Whilst other studies have also reported the age and sex breakdown of admissions and outcomes of particular species, the emphasis of this study was on longitudinal data for a range of diverse species and therefore did not focus at that level of detail. Future studies within the region and comparative studies between regions could focus on age and sex as factors contributing to admissions and outcomes of certain species or animal groups. This data can also be mined as a tool for general wildlife monitoring.

Lastly, many admissions were eliminated from our analysis. This included cases in which a single cause for admission could not be distinguished. Again, this appears to be common practise in this style of study, and authors have addressed this differently. For example, by combining all traumas, or by including an “unknown” or “other” category. We opted to include as many clearly delineated admission categories as possible, based on information given upon presentation that is clarified by veterinary examination. Some CFA frequently occur together, such as car strikes of the mother leading to orphaned young, which further confounds exact numbers in each category. We predict that in cases where more than one CFA may be evident, the animal had a lower chance for survival, as studies have shown that trauma severity increases mortality risk [47]. Our subset analysis of CFA before and after the changes to data capture methods at AZWH, showed that, by and large, the top six CFA have remained constant (Supporting Table 5), primarily affecting the overt signs of disease category, admissions for which increased dramatically following this change (Supporting Figure 3, Supporting Table 5). The main impact was thus on the proportion of admissions we could include in our final dataset due to a more complete reporting system. However, overall sample sizes were robust, and the main findings of this study were not impacted.

## Conclusion

From our retrospective longitudinal study of wildlife admissions to a WRC, it is clear that direct and indirect human-related factors are key drivers of morbidity and mortality of wildlife in Australia. Car strikes, entanglements and attacks by domestic pets accounted for over 80% of all admissions, and together these admission categories had low survival rates compared to “natural” causes for admission.

We observed a steady increase in the number of admissions to AZWH that mirrors the increasing human population in the corresponding area. Whilst we did not directly measure habitat-fragmentation and -loss in this study, its effects are evident and the continued population growth and consequential urban expansion in this area will inevitably be accompanied by land clearing and habitat modification. We predict that without intervention, this will result in a continued increase in admissions and ultimately, the ongoing decline of local wildlife populations.

Given the above, it stands to reason that substantial, human-driven conservation management is required to minimise the collateral damage wrought by modern civilisation. Hence, proactive and strategic management efforts to mitigate threats to biodiversity, and to the survival of wild populations of native species are an imminent and critical need, and it is also critical that these are underpinned by overarching legislative control and policy to balance the needs for human development alongside the conservation of biodiversity. Anthropogenic threats may be minimised by thoughtful landscape scale planning, incorporating biological corridors, strategic habitat restoration and defragmentation, as well as measures to minimise the spread of infectious diseases. Education, awareness and fundraising campaigns regarding thoughtful pet ownership alongside wildlife friendly driving habits and conservation strategies that aim to mitigate threats posed by feral animals will also be a step toward ameliorating the detrimental effects of human activities on wildlife. Without significant action, we are likely to are likely to see indelible changes to the unique Australian biota including more human-induced localised extinctions and the decline of species that are currently deemed ‘common’.

## Supporting information

### Supporting Files

**Supporting file 1:** List of species and species pools (sorted into animal groups) studied between 2006 and 2017.

**Supporting file 2:** List of causes for admission studied between 2006 and 2017.

### Supporting Tables

**Supporting Table 1:** Number of monthly admissions to AZWH per species or multi-species group between January 2006 and December 2017 (inclusive).

**Supporting Table 2:** Number of admissions to AZWH in each CFA category.

**Supporting Table 3:** Outcomes of the top six CFA. Raw values and proportions of admissions for each species or multi-species group are both presented.

**Supporting Table 4:** Odds ratio and relative risk analysis for the top six CFA, for each animal group.

**Supporting Table 5:** Analysis of changes to the order of the top six CFA following changes to database capture.

### Supporting Figures

**Supporting Figure 1:** Animal admissions to the Australia Zoo Wildlife Hospital between January 2006 and December 2017 (inclusive). Total annual (a) and average (b) admissions per animal group. Taxa are coloured based on higher classifications; see legend.

**Supporting Figure 2:** Monthly animal admissions to AZWH between January 2006 and December 2017 (inclusive) for each animal group: (a) avians; (b) reptiles; (c) amphibians; (d) marsupial mammals; (e) eutherian mammals. Trend lines are included to highlight the overall increase in admissions over the study period. Note the different Y axis ranges.

**Supporting Figure 3:** Monthly animal admissions to AZWH between January 2006 and December 2017 (inclusive) for the top six CFA: (a) hit by car; (b) overt signs of disease; (c) orphaned/dependent young; (d) entanglements; (e) dog attacks; (f) cat attacks. Trend lines are included to highlight the overall increase in admissions over the study period. Note the different Y axis ranges.

**Supporting Figure 4:** Seasonality of animal admissions to AZWH between January 2006 and December 2017 (inclusive) for the top six CFA: (a) hit by car; (b) overt signs of disease; (c) orphaned/dependent young; (d) entanglements; (e) dog attacks; (f) cat attacks. The mean per animal group is shown. Taxa are coloured based on higher classifications; see legend. Note the different Y axis ranges.

**Supporting Figure 5:** Outcomes of the top six CFA. Values depicted are the proportions of total admissions for each species or multi-species group, for each CFA: (a) hit by car; (b) overt signs of disease; (c) orphaned/dependent young; (d) entanglements; (e) dog attacks; (f) cat attacks. Taxa are ordered per mortality rate (beige bars); note the different order for graphs (a) to (f).

## Acknowledgements

We are greatly appreciative of input from the Australia Zoo Wildlife hospital, especially Kathy Whitefield. We highly value the continued contribution of volunteers and wildlife carers.

## Author contributions

ATB analysed and interpreted the data and wrote the manuscript. RB and AG assisted in data interpretation and wrote the manuscript. EM conducted database extraction and wrote the manuscript. RB, AG, SO, AP and GC conceived the study and wrote the manuscript. All authors reviewed the manuscript.

## References

1. Burton E, Tribe A. The Rescue and Rehabilitation of Koalas (*Phascolarctos cinereus*) in Southeast Queensland. Animals. 2016;6(9): 56.

2. Canfield PJ. A survey of koala road kills in New South Wales. Journal of Wildlife Diseases. 1991;27(4): 657–60.

3. Dique DS, Thompson J, Preece HJ, Penfold GC, de Villiers DL, Leslie RS. Koala mortality on roads in south-east Queensland: the koala speed-zone trial. Wildlife research. 2003;30(4): 419–26.

4. Gonzalez-Astudillo V, Allavena R, McKinnon A, Larkin R, Henning J. Decline causes of Koalas in South East Queensland, Australia: a 17-year retrospective study of mortality and morbidity. Scientific Reports. 2017;7:42587.

5. Griffith JE, Dhand NK, Krockenberger MB, Higgins DP. A retrospective study of admission trends of koalas to a rehabilitation facility over 30 years. J Wildl Dis. 2013;49(1): 18–28. doi: 10.7589/2012–05–135. PubMed PMID: 23307368.

6. Scheelings TF, Frith SE. Anthropogenic Factors Are the Major Cause of Hospital Admission of a Threatened Species, the Grey-Headed Flying Fox (*Pteropus poliocephalus*), in Victoria, Australia. PloS one. 2015;10(7):e0133638.

7. Scheelings TF. Morbidity and mortality of monotremes admitted to the Australian Wildlife Health Centre, Healesville Sanctuary, Australia, 2000–2014. Aust Vet J. 2016;94(4): 121–4. doi: 10.1111/avj.12417. PubMed PMID: 27021894.

8. Scheelings TF. Morbidity and Mortality of Reptiles Admitted to the Australian Wildlife Health Centre, Healesville Sanctuary, Australia, 2000–13. J Wildl Dis. 2015;51(3): 712–8. doi: 10.7589/2014–09–230. PubMed PMID: 26161722.

9. Woinarski JCZ, Burbidge AA, Harrison PL. Ongoing unraveling of a continental fauna: Decline and extinction of Australian mammals since European settlement. Proceedings of the National Academy of Sciences. 2015;112(15): 4531–40. doi: 10.1073/pnas.1417301112.

10. Andrén H, Linnell JD, Liberg O, Andersen R, Danell A, Karlsson J, et al. Survival rates and causes of mortality in Eurasian lynx (*Lynx lynx*) in multi-use landscapes. Biological Conservation. 2006;131(1): 23–32.

11. Erickson WP, Johnson GD, David Jr P. A summary and comparison of bird mortality from anthropogenic causes with an emphasis on collisions. 2005.

12. Ferreras P, Aldama J, Beltran J, Delibes M. Rates and causes of mortality in a fragmented population of Iberian lynx *Felis pardina Temminck*, 1824. Biological conservation. 1992;61(3): 197–202.

13. Molina-López R, Darwich L. Causes of admission of little owl (*Athene noctua*) at a wildlife rehabilitation centre in Catalonia (Spain) from 1995 to 2010. Animal Biodiversity and Conservation. 2011;34(2): 401–5.

14. Molina-López RA, Casal J, Darwich L. Causes of morbidity in wild raptor populations admitted at a wildlife rehabilitation centre in Spain from 1995–2007: a long term retrospective study. PLoS One. 2011;6(9):e24603.

15. Schmidt-Posthaus H, Breitenmoser-Wörsten C, Posthaus H, Bacciarini L, Breitenmoser U. Causes of mortality in reintroduced Eurasian lynx in Switzerland. Journal of Wildlife Diseases. 2002;38(1): 84–92.

16. Seiler A, Helldin J. Mortality in wildlife due to transportation. The ecology of transportation: managing mobility for the environment: Springer; 2006. p. 165–189.

17. Wendell MD, Sleeman JM, Kratz G. Retrospective study of morbidity and mortality of raptors admitted to Colorado State University Veterinary Teaching Hospital during 1995 to 1998. Journal of Wildlife Diseases. 2002;38(1): 101–6.

18. Brand CJ. Wildlife mortality investigation and disease research: contributions of the USGS National Wildlife Health Center to endangered species management and recovery. Ecohealth. 2013;10(4): 446–54. doi: 10.1007/s10393–013–0897–4. PubMed PMID: 24419670; PubMed Central PMCID: PMCPMC3938848.

19. Grogan A, Kelly A. A review of RSPCA research into wildlife rehabilitation. Vet Rec. 2013;172(8): 211. doi: 10.1136/vr.101139. PubMed PMID: 23436601.

20. Heigl F, Horvath K, Laaha G, Zaller JG. Amphibian and reptile road-kills on tertiary roads in relation to landscape structure: using a citizen science approach with open-access land cover data. BMC Ecol. 2017;17(1): 24. doi: 10.1186/s12898–017–0134-z. PubMed PMID: 28651557; PubMed Central PMCID: PMCPMC5485744.

21. Loss SR, Will T, Marra PP. The impact of free-ranging domestic cats on wildlife of the United States. Nature communications. 2013;4:1396.

22. McRuer DL, Gray LC, Horne L-A, Clark EE. Free-roaming cat interactions with wildlife admitted to a wildlife hospital. The Journal of Wildlife Management. 2017;81(1): 163–73. doi: 10.1002/jwmg.21181.

23. Montesdeoca N, Calabuig P, Corbera JA, Oros J. A long-term retrospective study on rehabilitation of seabirds in Gran Canaria Island, Spain (2003–2013). PLoS One. 2017;12(5):e0177366. doi: 10.1371/journal.pone.0177366. PubMed PMID: 28475653; PubMed Central PMCID: PMCPMC5419649.

24. Schenk AN, Souza MJ. Major anthropogenic causes for and outcomes of wild animal presentation to a wildlife clinic in East Tennessee, USA, 2000–2011. PLoS One. 2014;9(3):e93517. doi: 10.1371/journal.pone.0093517. PubMed PMID: 24686490; PubMed Central PMCID: PMCPMC3970955.

25. Tejera G, Rodriguez B, Armas C, Rodriguez A. Wildlife-vehicle collisions in Lanzarote Biosphere Reserve, Canary Islands. PLoS One. 2018;13(3):e0192731. doi: 10.1371/journal.pone.0192731. PubMed PMID: 29561864; PubMed Central PMCID: PMCPMC5862401.

26. Morner T. Health Monitoring and Conservation of Wildlife in Sweden and Northern Europe. Annals of the New York Academy of Science. 2002;969:34–8.

27. United Nations. World Population Prospects: The 2017 Revision, Key Findings and Advance Tables. In: Department of Economic and Social Affairs PD, editor. New York: Working Paper No. ESA/P/WP/248; 2017.

28. Ceballos G, Ehrlich PR. Mammal population losses and the extinction crisis. Science. 2002;296(5569): 904–7.

29. Ceballos G, García A, Ehrlich PR. The sixth extinction crisis: loss of animal populations and species. Journal of Cosmology. 2010;8(1821):31.

30. Barnosky AD, Matzke N, Tomiya S, Wogan GO, Swartz B, Quental TB, et al. Has the Earth/’s sixth mass extinction already arrived? Nature. 2011;471(7336): 51–7.

31. Lewis OT. Climate change, species–area curves and the extinction crisis. Philosophical Transactions of the Royal Society of London B: Biological Sciences. 2006;361(1465):163–71.

32. Ceballos G, Ehrlich PR, Barnosky AD, García A, Pringle RM, Palmer TM. Accelerated modern human–induced species losses: Entering the sixth mass extinction. Science advances. 2015;1(5):e1400253.

33. Lowry H, Lill A, Wong BB. Behavioural responses of wildlife to urban environments. Biol Rev Camb Philos Soc. 2013;88(3): 537–49. doi: 10.1111/brv.12012. PubMed PMID: 23279382.

34. McKinney ML. Urbanization, Biodiversity, and Conservation: The impacts of urbanization on native species are poorly studied, but educating a highly urbanized human population about these impacts can greatly improve species conservation in all ecosystems. BioScience. 2002;52(10): 883–90. doi: doi.org/10.1641/0006–3568(2002)052[0883:UBAC]2.0.CO;2.

35. McDonald R, Marcotullio P. Global Effects of Urbanization on Ecosystem Services. In: Niemelä J, Breuste JH, Elmqvist T, Guntenspergen G, James P, McIntyre NE, editors. Urban Ecology: Patterns, Processes, and Applications: Oxford Scholarship Online; 2011.

36. Lai S, Loke LHL, Hilton MJ, Bouma TJ, Todd PA. The effects of urbanisation on coastal habitats and the potential for ecological engineering: A Singapore case study. Ocean Coast Manage. 2015;103:78–85. doi: 10.1016/j.ocecoaman.2014.11.006. PubMed PMID: WOS:000347769600009.

37. Finn HC, Stephens NS. The invisible harm: land clearing is an issue of animal welfare. Wildlife Research. 2017;44(5): 377. doi: 10.1071/wr17018.

38. Gebreab SZ, Vienneau D, Feigenwinter C, Ba H, Cisse G, Tsai MY. Spatial air pollution modelling for a West-African town. Geospat Health. 2015;10(2): 321. doi: 10.4081/gh.2015.321. PubMed PMID: 26618306.

39. Moiron M, González-Lagos C, Slabbekoorn H, Sol D. Singing in the city: high song frequencies are no guarantee for urban success in birds. Behavioural Ecology. 2015;26(3): 843–50. doi: doi.org/10.1093/beheco/arv026.

40. Gaston KJ, Visser ME, Hölker F. The biological impacts of artificial light at night: the research challenge. Philosophical Transactions of the Royal Society B. 2015;370(1667). doi: 10.1098/rstb.2014.0133.

41. Andersson MN, Wang H-L, Nord A, Salmón P, Isaksson C. Composition of physiologically important fatty acids in great tits differs between urban and rural populations on a seasonal basis. Frontiers in Ecology and Evolution. 2015;3(93). doi: 10.3389/fevo.2015.00093.

42. Isaksson C. Urbanization, oxidative stress and inflammation: a question of evolving, acclimatizing or coping with urban environmental stress. Functional Ecology. 2015;29(7): 913–23. doi: 10.1111/1365–2435.12477

43. Brahmia Z, Scheifler R, Crini N, Maas S, Giraudoux P, Benyacoub S. Breeding performance of blue tits (*Cyanistes caeruleus ultramarinus*) in relation to lead pollution and nest failure rates in rural, intermediate, and urban sites in Algeria. Environ Pollut. 2013;174:171–8. doi: 10.1016/j.envpol.2012.11.028. PubMed PMID: 23262073.

44. Sumasgutner P, Nemeth E, Tebb G, Krenn HW, Gamauf A. Hard times in the city - attractive nest sites but insufficient food supply lead to low reproduction rates in a bird of prey. Front Zool. 2014;11:48. doi: 10.1186/1742–9994–11–48. PubMed PMID: 24872836; PubMed Central PMCID: PMCPMC4035672.

45. Steyn L, Maina JN. Comparison of the numbers of free (surface) macrophages in the respiratory systems of three species of birds in an urban and a rural area of South Africa. Journal of Ornithology. 2015;156(4): 1085–93.

46. Salafsky N, Salzer D, Stattersfield AJ, HILTON-TAYLOR C, Neugarten R, Butchart SH, et al. A standard lexicon for biodiversity conservation: unified classifications of threats and actions. Conservation Biology. 2008;22(4): 897–911.

47. Mullineaux E. Veterinary treatment and rehabilitation of indigenous wildlife. J Small Anim Pract. 2014;55(6): 293–300. doi: 10.1111/jsap.12213. PubMed PMID: 24725160.

48. Pyke GH, Szabo JK. Conservation and the 4 Rs, which are rescue, rehabilitation, release, and research. Conserv Biol. 2018;32(1): 50–9. doi: 10.1111/cobi.12937. PubMed PMID: 28328146.

49. Bouchon-Small A. The rescue and rehabilitation of wildlife in South East Queensland with a case study of the birds of prey: University of Queensland; 2015 (Thesis).

50. Martínez JC, Rosique AI, Royo MS. Causes of admission and final dispositions of hedgehogs admitted to three Wildlife Rehabilitation Centers in eastern Spain. Hystrix, the Italian Journal of Mammalogy. 2014;25(2): 107–10. doi: 10.4404/hystrix-25.2–10248.

51. Hamer AJ, McDonnell MJ. The response of herpetofauna to urbanization: Inferring patterns of persistence from wildlife databases. Austral Ecology. 2010;35(5): 568–80. doi: doi:10.1111/j.1442–9993.2009.02068.x.

52. Pyke GH, Szabo JK. What can we learn from untapped wildlife rescue databases? The masked lapwing as a case study. Pacific Conservation Biology. 2018;24(2): 148. doi: 10.1071/pc18003.

53. Sleeman J. Use of wildlife rehabilitation centers as monitors of ecosystem health. Zoo and wild animal medicine. 2008;6:97–104.

54. Trocini S, Pacioni C, Warren K, Butcher J, Robertson I. Wildlife disease passive surveillance: the potential role of wildlife rehabilitation centres. National Wildlife Rehabilitation Conference; Canberra, ACT, Australia 2008.

55. Molina-Lopez RA, Darwich L. Causes of admission of little owl (*Athene noctua*) at a wildlife rehabilitation centre in Catalonia (Spain) from 1995 to 2010. Animal Biodiversity and Conservation. 2011;34(2): 401–5.

56. Rodríguez B, Rodríguez A, Siverio F, Siverio M. Causes of Raptor Admissions to a Wildlife Rehabilitation Center in Tenerife (Canary Islands). Journal of Raptor Research. 2009;44(1): 30–9. doi: doi.org/10.3356/JRR-09–40.1.

57. Santos SM, Lourenco R, Mira A, Beja P. Relative effects of road risk, habitat suitability, and connectivity on wildlife roadkills: the case of tawny owls (*Strix aluco*). PLoS One. 2013;8(11):e79967. doi: 10.1371/journal.pone.0079967. PubMed PMID: 24278226; PubMed Central PMCID: PMCPMC3836987.

58. Burton DL, Dobar KA, editors. Morbidity and mortality of urban wildlife in the midwestern United States. Proceedings 4th International Urban Wildlife Symposium; 2004.

59. ISO. International Organization for Standardization. Persistent stored modules (SQL/PSM). SQL2016.

60. IBM Corp. IBM SPSS Statistics for Windows, Version 24.0. Armonk, NY: IBM Corp.; 2016.

61. RSPCA: Wildlife Hospital; What we do 2017 [22/05/2018]. Available from: https://www.rspcaqld.org.au/what-we-do/care-for-animals/wildlife-hospital

62. Currumbin Wildlife Hospital Foundation: Wildlife Hospital 2018 [22/05/2018]. Available from: http://cwhf.org.au/wildlife-hospital/.

63. Wimberger K, Downs CT. Annual intake trends of a large urban animal rehabilitation centre in South Africa: a case study. Animal Welfare. 2010;19:501–13.

64. Molina-Lopez RA, Casal J, Darwich L. Causes of morbidity in wild raptor populations admitted at a wildlife rehabilitation centre in Spain from 1995–2007: a long term retrospective study. PLoS One. 2011;6(9):e24603. doi: 10.1371/journal.pone.0024603. PubMed PMID: 21966362; PubMed Central PMCID: PMCPMC3179465.

65. da Rosa CA, Bager A. Seasonality and habitat types affect roadkill of neotropical birds. J Environ Manage. 2012;97:1–5. doi: 10.1016/j.jenvman.2011.11.004. PubMed PMID: 22325576.

66. Census update: Sunshine Coast. In: Housing ABoSCoPa, editor. 2016.

67. Sunshine Coast Business Council: Sunshine Coast Region [Internet]. 2012 [cited 22/05/2018]. Available from: https://scbusinesscouncil.com.au/sunshine-coast-region/

68. Woinarski J, Burbidge AA. Phascolarctos cinereus. The IUCN Red List of Threatened Species 2016. In: IUCN, editor. 2016.

69. Beyer HL, Villiers Dd, Loader J, Robbins A, Stigner M, Forbes N, et al. Management of multiple threats achieves meaningful koala conservation outcomes. Journal of Applied Ecology. 2018;1(10). doi: DOI:10.1111/1365–2664.13127.

70. Devereaux LN, Polkinghorne A, Meijer A, Timms P. Molecular evidence for novel chlamydial infections in the koala (*Phascolarctos cinereus*). Systematic and applied microbiology. 2003;26(2): 245–53. doi: 10.1078/072320203322346092. PubMed PMID: 12866851.

71. Polkinghorne A, Hanger J, Timms P. Recent advances in understanding the biology, epidemiology and control of chlamydial infections in koalas. Vet Microbiol. 2013;165(3-4):214–23. doi: 10.1016/j.vetmic.2013.02.026. PubMed PMID: 23523170.

72. Brunton EA, Srivastava SK, Schoeman DS, Burnett S. Quantifying trends and predictors of decline in eastern grey kangaroo (*Macropus giganteus*) populations in a rapidly urbanising landscape. Pacific Conservation Biology. 2018;24(1): 63–73. doi: https://doi.org/10.1071/PC17034.

73. Ellis WA, FitzGibbon SI, Barth BJ, Niehaus AC, David GK, Taylor BD, et al. Daylight saving time can decrease the frequency of wildlife-vehicle collisions. Biol Lett. 2016;12(11). doi: 10.1098/rsbl.2016.0632. PubMed PMID: 27881767; PubMed Central PMCID: PMCPMC5134043.

74. Birds in Backyards. Autumn Survey - 2018 results: BirdLife Australia; 2018 [16/07/2018]. Available from: http://www.birdsinbackyards.net/content/article/Autumn-Survey-2018-results.

75. Pest & Disease Information Service. Rainbow lorikeet: management. WA Department of Primary Industries and Regional Development. South Perth, WA, Australia 2018.

76. Raidal SR, Sarker S, Peters A. Review of psittacine beak and feather disease and its effect on Australian endangered species. Aust Vet J. 2015;93(12): 466–70. doi: 10.1111/avj.12388. PubMed PMID: 26769072.

77. McOrist S, Reece RL. Clostridial enteritis in free-living lorikeets (*Trichoglossus spp*.). Avian Pathol. 1992;21(3): 503–7. doi: 10.1080/03079459208418868. PubMed PMID: 18670965.

78. NSW Government. The danger of feeding lorikeets. 2018.

79. Queensland Land for Wildlife. Wildlife Friendly Fencing and Netting. Healthy Land and Water; 2017.

80. Burbidge AA, Abbott I. Mammals on Western Australian islands: occurrence and preliminary analysis. Australian Journal of Zoology. 2017;65(3): 183–95. doi: https://doi.org/10.1071/ZO17046.

81. Woinarski JCZ, Legge S, Fitzsimons JA, Traill BJ, Burbidge AA, Fisher A, et al. The disappearing mammal fauna of northern Australia: context, cause, and response. Conservation Letters. 2011;4(3): 192–201. doi: doi:10.1111/j.1755–263X.2011.00164.x.

82. Dickman CR. House cats as predators in the Australian environment: impacts and management. Human–Wildlife Conflicts 2009;3(1): 41–8.

83. Abbott I. Origin and spread of the cat, *Felis catus* on mainland Australia, with a discussion of the magnitude of its early impact on native fauna. Wildlife Research. 2002;29(1): 51–74. doi: https://doi.org/10.1071/WR01011.

84. Dickman C. Overview of the impacts of feral cats on australian native fauna. University of Sydney Institute of Wildlife Research and School of Biological Sciences for the Australian Nature Conservation Agency; 1996.

85. Ancillotto L, Serangeli MT, Russo D. Curiosity killed the bat: Domestic cats as bat predators. Mamm Biol. 2013;78(5): 369–73. doi: 10.1016/j.mambio.2013.01.003. PubMed PMID: WOS:000323534000010.

86. Muhldorfer K, Speck S, Wibbelt G. Diseases in free-ranging bats from Germany. BMC Vet Res. 2011;7:61. doi: 10.1186/1746–6148–7–61. PubMed PMID: 22008235; PubMed Central PMCID: PMCPMC3219556.

87. Paltridge R. The diets of cats, foxes and dingoes in relation to prey availability in the Tanami Desert, Northern Territory. Wildlife Research. 2002;29(4): 389–403. doi: https://doi.org/10.1071/WR00010.

88. Risbey DA, Calver MC, Short J, Bradley JS, Wright IW. The impact of cats and foxes on the small vertebrate fauna of Heirisson Prong, Western Australia. II. A field experiment. Wildlife Research. 2000;27(3): 223–35. doi: https://doi.org/10.1071/WR98092.

89. Denny E, Dickman CR. Review of cat ecology and management strategies in Australia. Invasive Animals Cooperative Research Centre. Canberra, Australia:; 2010.

90. Letnic M, Koch F, Gordon C, Crowther MS, Dickman CR. Keystone effects of an alien top-predator stem extinctions of native mammals. Proc Biol Sci. 2009;276(1671):3249–56. doi: 10.1098/rspb.2009.0574. PubMed PMID: 19535372; PubMed Central PMCID: PMCPMC2817164.

91. Johnson CN, Isaac JL, Fisher DO. Rarity of a top predator triggers continent-wide collapse of mammal prey: dingoes and marsupials in Australia. Proc Biol Sci. 2007;274(1608):341–6. doi: 10.1098/rspb.2006.3711. PubMed PMID: 17164197; PubMed Central PMCID: PMCPMC1702374.

92. Glen AS, Dickman CR, Soulé ME, Mackey BG. Evaluating the role of the dingo as a trophic regulator in Australian ecosystems. Austral Ecology. 2007;32(5): 492–501. doi: doi:10.1111/j.1442–9993.2007.01721.x.

93. Letnic M, Ritchie EG, Dickman CR. Top predators as biodiversity regulators: the dingo *Canis lupus* dingo as a case study. Biological Reviews. 2012;87(2): 390–413. doi: doi:10.1111/j.1469–185X.2011.00203.x.

94. Cook TC, Blumstein DT. The omnivore’s dilemma: Diet explains variation in vulnerability to vehicle collision mortality. Biological Conservation. 2013;167:310–5. doi: https://doi.org/10.1016/j.biocon.2013.08.016.

95. Brown JD, Sleeman JM. Morbidity and mortality of reptiles admitted to the Wildlife Center of Virginia, 1991 to 2000. Journal of wildlife diseases. 2002;38(4): 699–705. doi: 10.7589/0090–3558–38.4.699. PubMed PMID: 12528435.

96. Bereau of Meteorology. Queensland in December 2010: The wettest December on record 2011 [16/05/2018]. Available from: http://www.bom.gov.au/climate/current/month/qld/archive/201012.summary.shtml.

97. Bureau of Meteorology. Brisbane in November 2016: Very warm days; dry 2016 [16/05/2018]. Available from: http://www.bom.gov.au/climate/current/month/qld/archive/201611.brisbane.shtml.

98. Bureau of Meteorology. Queensland in spring 2014: Dry over large parts of the State; some extremely hot days 2014 [16/05/2018]. Available from: http://www.bom.gov.au/climate/current/season/qld/archive/201411.summary.shtml.

99. Bureau of Meteorlogy. Brisbane in November 2014: A hot month; severe thunderstorms lash the city 2014 [16/05/2018]. Available from: http://www.bom.gov.au/climate/current/month/qld/archive/201411.brisbane.shtml.

100. Bureau of Meteorology. Annual climate statement 2017. 2018 [16/05/2018]. Available from: http://www.bom.gov.au/climate/current/annual/aus/.

101. Mullineaux E, Kidner P. Managing public demand for badger rehabilitation in an area of England with endemic tuberculosis. Vet Microbiol. 2011;151(1–2):205–8. doi: 10.1016/j.vetmic.2011.02.045. PubMed PMID: WOS:000292352700029.

102. Kelly A, Bland M. Admissions, diagnoses, and outcomes for Eurasian Sparrowhawks (*Accipiter nisus*) brought to a wildlife rehabilitation center in England. Journal of Raptor Research. 2006;40(3): 231–5. doi: Doi10.3356/0892–1016(2006)40[231:Adaofe]2.0.Co;2. PubMed PMID: WOS:000241997700009.

